# Nicotine causes highly localized and tuned changes in thalamocortical gain that cascade strongly and selectively through the cortical circuit

**DOI:** 10.64898/2026.02.22.707315

**Authors:** Veronica C. Galvin, Anita A. Disney

## Abstract

In layer 4 of the primary sensory cortices, sensory data meets contextual information carried by modulatory systems, establishing a potentially important point of control. Accordingly, modulatory receptor expression at this point often differs from other nearby layers and cortical areas. Altering the processing state at even a small number of layer 4 thalamocortical synapses should be impactful on downstream processing, as this is the point at which most information about the outside world enters cortex. It is, however, often assumed—based on diffuse innervation of cortex by subcortical nuclei, and dense receptor expression in the association cortices—that modulatory signals are shared across large swaths of tissue, with the primary site of action being in ‘higher’ levels of cortex. Hypothesizing that modifying the feedforward input to cortex might be an important capability of the cholinergic system, we recorded across the depth of cortex while delivering nicotine to a small proportion of thalamocortical synapses in macaque primary visual cortex, leveraging selective receptor expression to achieve spatial control. We observed bidirectional response changes throughout all layers, including layers in which nicotine is known to have no effect when applied locally. The pattern of these changes carried the signature of having passed through a normalizing circuit, rather than signatures of drug spread: the magnitude and tuning dependence of the response changes were well-accounted for by a normalization model with a tuned, multiplicative gain field. The circuit apparently did not compensate these changes; perceived contrast in a behavioral task was biased with the same normalization-predicted tuning dependence.

**Significance:** In seeking to understand the role(s) neuromodulators play in cognition and behavior, answers are generally sought in the circuits of the association cortices, and neuromodulation is often conceived of, and modelled, in a manner akin to a coarse-grained—even cortex-wide—volume knob. Here, we show that focal cholinergic modulation of thalamocortical transmission in the visual system reshapes columnar neural activity and alters perceptual performance. Our observations suggest an underappreciated role for an old mechanistic concept: modulatory control of cortical gating.

## Introduction

The convergence, in layer 4 of cortex, of pathways carrying sensory data with modulatory systems carrying state information (Prusky et al., 1987; Rakic et al., 1988; Watakabe et al., 2009; Zilles and Palomero-Gallagher, 2017 ; Disney, 2021 ; Rapan et al., 2022) yields an important architecture for ‘gating’ control (Moruzzi and Magoun, 1949). What propagates through layer 4—and in what form—constrains representations across the rest of the cortex. Under the implied control this anatomy suggests, it would be difficult to know to what extent deficits observed when neuromodulatory signalling is disrupted or disordered arise from failed cognitive processing versus perfectly good cognition attempting to operate on flawed data, or some combination of the two.

Given the important gating role layer 4, and thalamocortical synapses specifically, play in the flow of data to the circuits of cortex, control of information passing through even a small proportion of these synapses should impact downstream processing. Importantly, our assertion here is not that adding or mimicking sensory data at this point in the circuit is impactful—this seems obviously true; we can perceive small and/or faint stimuli.

Accordingly, causal methods that alter spiking irrespective of sensory data—such as electrical and optogenetic stimulation—can reorganize circuit activity and alter perception and behavior (Murphey and Maunsell, 2007; Jazayeri et al., 2012; Tehovnik and Slocum, 2013; De et al., 2020). Instead, our hypothesis is that neuromodulatory control of only the *processing state* of a thalamocortical synapse should importantly alter downstream activity.

Testing this hypothesis requires that four conditions be met: We need 1) a circuit in which a modulatory receptor is uniquely expressed at the thalamocortical synapse (and nowhere else nearby), 2) a selective ligand for that receptor, 3) a means to parametrically vary the strength of sensory drive, and 4) a means to read out gain at the neural and behavioral level. In macaque primary visual cortex (V1), cholinergic β2 subunit-containing nicotinic acetylcholine receptors (nAChRs) are expressed presynaptically on most thalamocortical synapses onto excitatory neurons in the primary thalamic recipient sublayer, 4C; they are only very sparsely expressed elsewhere in V1. Activating these receptors does not alter activity without a visual stimulus, but yields a profound, multiplicative increase on visual responses (Disney et al., 2007).

Our framing implies that a modulatory system with receptors expressed at the gate to cortex—as is the case for the cholinergic system—should have the capacity to profoundly impact processing. Indeed, lesions of the basal forebrain (whose afferents deliver all acetylcholine to cortex in non-rodent species) eliminate visual responses for most V1 neurons (Sato et al., 1987). However, local delivery of nicotine in macaque V1, <u>without</u> <u>targeting layer 4C</u>, does not alter visual responses (Disney et al., 2007) or performance on a visuospatial attention task (Herrero et al., 2008). In fact, to our knowledge, only delivery of large volumes (via microinfusion or systemic delivery) of any drug targeting neuromodulation, in any region of macaque cortex, has been shown to alter behavior (Sawaguchi and Goldman-Rakic, 1991; Li et al., 1999; Vijayraghavan et al., 2007; Wang et al., 2007; Herrero et al., 2008; Noudoost and Moore, 2011b; Thiele et al., 2012; Seillier et al., 2017; Vijayraghavan et al., 2018; Dasilva et al., 2019; Herrero and Thiele, 2021). Numerous studies report changes in <u>neural responses</u> when small volumes of modulatory drugs are delivered. Do they just peter out? Or are they actively compensated? And: Would changes applied at thalamocortical synapses be different?

We delivered nanoliter volumes of nicotine to layer 4C while recording neural activity across the full depth of the V1 column. We found that: 1) in layer 4C, only visually driven activity changed; 2) effects propagated to yield bidirectional changes in responses across V1 all layers that were; 3) well-captured by a normalization model of attention with a tuned gain field. Finally, 4) this focal gain modulation of thalamocortical transmission by nicotine altered perceptual performance.

## Methods

### Animals

One adult male (D, age 10) and one adult female (E, age 12) *Macaca mulatta* were used in this study. Animals were socially housed in an AAALAC-certified facility at Duke University on a 12-hour light/dark cycle in humidity and temperature-controlled environments. All procedures were approved by Duke University’s Institutional Animal Care and Use Committee, in accordance with NIH guidelines. Both animals were maintained on *ad libitum* food and water in their home cages throughout the study.

### Behavior

*Behavioral control and monitoring*: During task performance and recording, head fixed (Thomas PPS ‘halo’ system) animals were seated in a dimly lit (< 1 cd/m^2^) Faraday room and monitored continuously via infrared camera. Eye position and pupil size were monitored non-invasively using a high-speed infrared eye tracking system (EyeLink). Visual stimuli were delivered by a ProPixx projector (VPixx) onto the central 73.5 x 40.5 cm of a 122 x 91 cm (W x H) polarization-preserving rear projection screen (Stewart Filmscreen) positioned 98.5 cm from the eyes.

*Passive Fixation Task:* Animals held gaze within 1.0 degree of visual angle (dva) of a fixation point at the center of the screen while Gabor stimuli of varying position, orientation, and contrast (diameter = 2 dva, spatial frequency = 1.5 cycles/degree, cpd) were presented for 150-ms on/off periods on a 20 cd/m^2^ mean gray background (**Fig. 1A**). Accurate fixation was rewarded with ∼0.2 mL dilute juice. In one animal (D) the Gabor stimuli were static; in the other (E) they were drifting.

**Figure 1.**
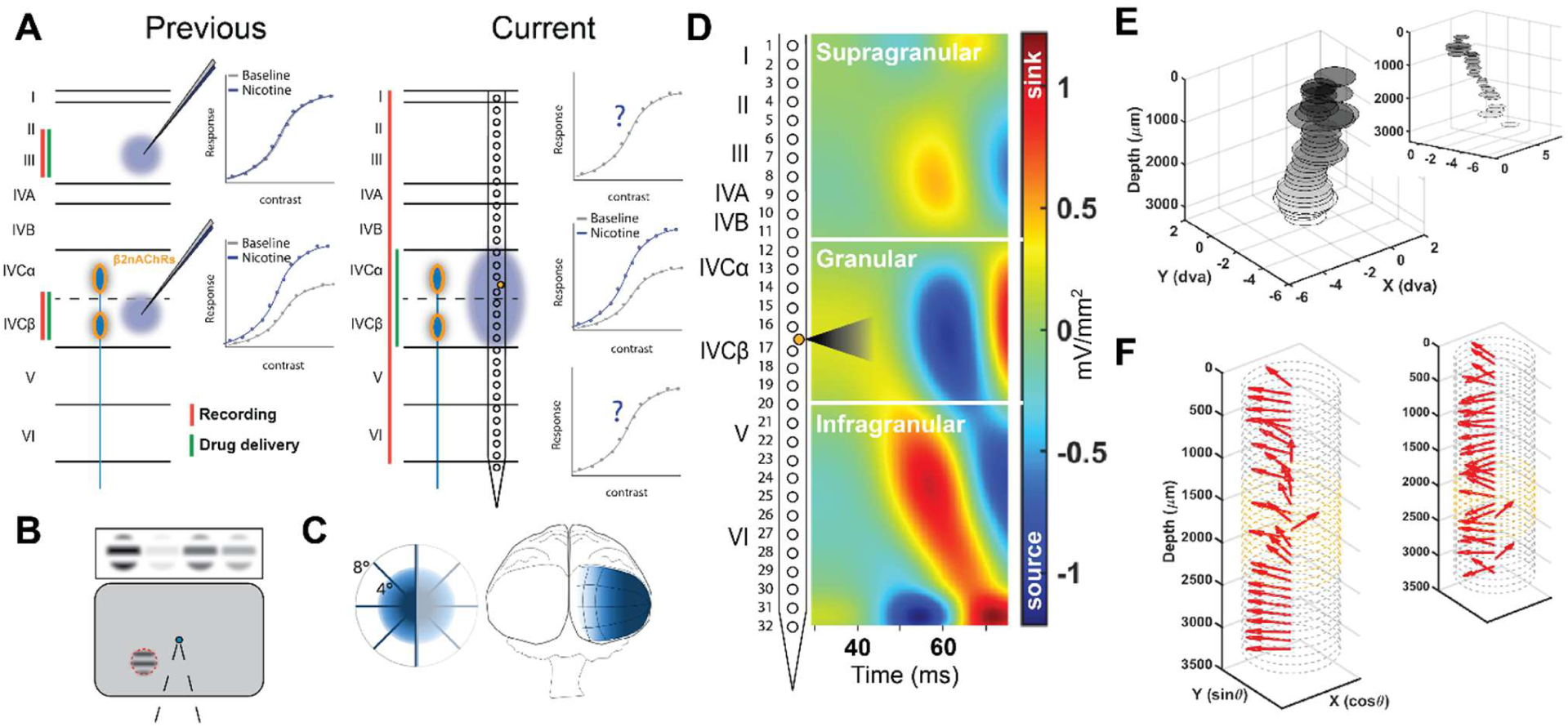
Experimental Design. **(A)** Prior studies that have combined local delivery of pharmacological agents (including nicotine) while recoding neural responses, have applied the drug (blue cloud and green vertical bar) and recorded unit activity (red vertical bar) in the same small volume of tissue (left: ‘Previous’). In this study (right: ‘Current’) recorded units can be up to 1.5 mm away from the site of drug delivery, in the middle of the electrode. Drug solutions are known to wick along much, or all, of the shank of these electrodes, the restriction of the causal manipulation to layer 4C relies on restricted nicotinic receptor expression (orange highlight on schematized blue thalamic axon arriving in layer 4C), not on port position. **(B)** Task schematic: Monkeys fixate centrally (blue dot; accuracy window 1° visual angle, dva) while Gabor stimuli are used to measure receptive field position (red dashed circle), orientation tuning, and contrast responses. **(C)** Recording sites were in parafoveal (<8° eccentricity) V1. **(D)** Electrode arrays were inserted roughly perpendicular to the brain surface and spanned the full cortical depth (left). A fluid capillary exit on the array (orange circle) was positioned close to layer 4C by current source density analysis of visually-evoked local field potentials and nicotine delivered by pressure ejection (black triangle). **(E)** Receptive field alignment across the 32 concurrently-recorded channels for our most (main) and least (inset) aligned (with the cortical column) recording days. Depth axis: 0 = ∼pia; 3200 μm = channel 32. Position: dva from fixation. **(F)** Preferred orientation (red arrows; circles: 360° of direction tuning) at each channel for the same most- and least-aligned days shown in D. Yellow dashed circles: layer 4C.

*Perceptual Equivalence Task:* After a fixation period of 750-1750 ms (1 dva accuracy), two static Gabor stimuli appeared simultaneously, one in each hemifield, equidistant from the fixation point. Stimuli were identical in size, spatial frequency, and orientation, but differed in contrast. Trials were randomly interleaved for two series of contrast comparisons against a) 20% and b) 40% contrast ‘reference’ Gabors. Stimuli for comparison were generated representing three half-steps of contrast for each reference and were presented in randomized order using the method of constant stimuli. The task was to saccade to the higher contrast stimulus. A juice reward was delivered for correct choices; choices between identical stimuli were randomly rewarded. To prevent reward feedback interfering with nicotine effects, during nicotine delivery a probabilistic reward was enacted such that stimuli that differed by 3% contrast or less were rewarded with probability 0.75, regardless of choice.

### Behavioral Analysis

The effect of nicotine did not depend on contrast pedestal, so performance was analyzed by combining trials across the two reference contrasts (20% and 40%), after normalizing to the reference (yielding contrast ratios: 0, 0.5, 0.75, 1.0, 1.25, 1.5, and 2.0). For each session, drug condition, and hemifield, we computed the proportion of trials on which the animal chose the comparison stimulus at each contrast ratio. Logistic functions of the form:

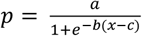

were fit to these curves using nonlinear least-squares regression (MATLAB Curve Fitting Toolbox), and the point of subjective equivalence (PSE) extracted as the contrast ratio yielding p = 0.5. We found no difference in behavioral performance between baseline (no delivery) and saline conditions, and so these trials were combined for comparison with nicotine.

We computed a composite score for each session capturing the net change in side-bias under nicotine. For each hemifield, we calculated the mean proportion chosen under nicotine and under saline. The composite was defined as (right_nicotine − right_saline) − (left_nicotine − left_saline), representing the net shift in rightward bias. This composite was then multiplied by +1 for co-tuned sessions (where nicotine, having been delivered to left V1 in both animals, is predicted to increase right hemifield apparent contrast) and by −1 for non-co-tuned sessions (where nicotine is predicted to decrease it). ‘Co-tuned’ was defined based on the range of gain field orientation tuning bandwidths fit to out of sample (neural) data using the normalization model below. A positive direction-corrected composite indicates a shift in the predicted direction. Statistical significance was assessed using a two-tailed binomial sign test and a one-sample t-test against zero.

### Electrophysiology

The combined electrophysiology and nicotine delivery experiments were conducted using 32-channel S-Probes with electrode spacing at 100 µm pitch, and a fluid capillary between electrodes 16 and 17 (**Fig. 1C**). Data were recorded using Synapse software from Tucker Davis Technologies (TDT) at a sampling rate of ∼24,414 kHz, amplified using the TDT PZ5 and RZ2 bioamp processors, band pass filtered (0.3 to 10 kHz), and stored for offline analysis.

S-Probes were positioned using a motorized drive (Thomas Recording), with a sharpened stainless steel guide tube to assist in passing dura. Using a single unit electrode (FHC), dura thickness was measured before the first recordings were made, and the approximate daily thickness estimated based on changes in height from the chamber top. Using these estimates, guide tube tips were positioned just before cortex (i.e., still within dura) and the stainless steel-tipped S-probe used to exit dura and enter cortex. Electrodes were advanced at a drive speed of 2 μm/second until the uppermost channel was just above, or within, layer 1 and allowed to rest in place for 20-60 minutes before positioning with respect to the granular layer was assessed by current source density (CSD) analysis. CSDs were calculated from a ∼2-minute recordings of local field potential responses to 100 ms flashes of 100% contrast, full-field gratings (1.5 cpd). Electrode position was adjusted as needed to place the fluid capillary exit (‘fluid port’) near the granular layer CSD sink (**Fig. 1C**); an additional 15-45min rest period was allowed after any depth adjustments, and the CSD result re-checked. Recorded units were assigned to layers (granular/4C, supragranular, infragranular) based on CSD plots.

Receptive field x-y position was mapped using flashed Gabor stimuli (2 dva, 1.5 cpd, 96% contrast, positions randomly interleaved). Orientation/direction tuning preference was measured in 22.5° steps (randomly interleaved, including blanks). In one animal orientation tuning was obtained (static Gabors; Monkey D); direction tuning in the other (drifting Gabors; Monkey E). Based on receptive field positions and tuning preferences across the S-Probe array, a ‘best-fit’ stimulus (x-y position and orientation/direction tuning) was chosen for use in measuring the contrast response function gain. Contrast responses were measured in 9 logarithmic steps from 1% to 96%, plus 0% contrast ‘blanks’ (all randomly interleaved).

### Pharmacology

Nicotine (Glentham Life Sciences; cat#65-31-6) was dissolved in sterile saline, pH adjusted to 4.5 - 7.0, and filtered at 0.22 µm prior to loading into electrodes for use. Pre-mixed solution was kept at 4°C for up to 7 days. Nicotine concentration differed across days (200 - 500 mM). At the start of each recording day the electrode fluid capillary was filled with saline and connected to a pneumatic pump via microtubing filled with nicotine solution.

One pulse of solution (saline or nicotine) was delivered by pressure ejection (WPI PicoPump PV830 or Medical Systems Corp PLI_100) between the onset of fixation and presentation of the first Gabor stimulus. An electrode-specific pressure range that delivered a droplet volume of ∼1-5 nL was determined *ex vivo* based on calibration under a microscope (volume estimation from measured droplet size). Ejection pressure was selected within this range, per-session, based on response stationarity (qualitatively assessed) for electrodes flanking the fluid port; final pressures ranged from 2.5-20 PSI. The number of possible saline control trials was constrained by the estimated number of delivery pulses to clear the electrode of saline. After the saline controls, we employed a 1-10-minute priming period of intermittent pulse delivery with no visual stimulation or behavioral control. Between 2 and 3 nicotine conditions were then collected at differing backing pressures, always from lowest to highest. The maximum nicotine ‘dose’ was the backing pressure matched to the highest-pressure saline condition that yielded stable recording. Either two (∼30-50% and 100% of the maximum dose) or three (33%, 66%, and 100% doses) were tested. The total volume of nicotine delivered for a given day (all dose levels, all trials, over 2-3 hours) was ∼500 - 1000 nL. The corresponding total volume of saline delivered per recording day was ∼300 – 500 nL, for a total volume (all solutions) of < 2 µL.

### Data Processing and Analysis

#### Spike recording, sorting, and windowing

Waveforms were sorted offline in MATLAB (MathWorks) using code developed by Dr. K. Nielsen (available on the nielsenlab GitHub repository; Khamiss et al., 2026). Thresholds were set per channel and sorted based on multiple waveform characteristics (spike amplitude maximum, minimum, and spike width). Response window start times were calculated per trial and per unit individually by calculating the average and standard deviation (sd) of the spike rate in the first 20 ms after stimulus onset, then finding the first 10 ms window during which the average rate was >4 sd over baseline. Response window length was 150 ms.

#### Inclusion criteria for analysis

Permissive inclusion criteria were used: units needed to have (1) a clear receptive field position, (2) a spike rate difference > 1 spike/sec between the peak and trough of the orientation tuning fit, (3) an average spiking response to the ‘best fit’ 96% contrast stimulus at least 1 sd higher than the average response to the 0% contrast blank, (4) an average response to the ‘best fit’ 96% contrast stimulus under the no-fluid delivery baseline of > 1 spike/sec, and (5) (for supragranular and infragranular only) a response that was not significantly suppressed by saline delivery, determined by a bootstrap analysis on the normalized response area (see Non-parametric analysis). These criteria are deliberately permissive: restricting to tightly tuned units would bias the sample toward those the model described below is best able to predict. Analyses restricted to well-driven units, and sensitivity analyses on inclusion criteria are described in the Results, below. Receptive field size and position were fit with a 2D gaussian function and double von Mises fits were applied to orientation/direction tuning.

#### Parametric analysis of contrast responses

A hyperbolic ratio (Naka-Rushton) function was fit for each condition (baseline, saline, drug) to the averaged (across 39-247 repeats) responses, R, for each stimulus contrast, C:

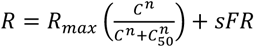

This fit yields four parameters: Rmax (maximum response), c50 (contrast at the half-maximum response), n (function slope), and sFR (offset attributable to ‘spontaneous’ firing).

To classify gain types (contrast gain, response gain, mixed, no gain change), we used bootstrap fits computed from 1000 iterations of resampling with replacement from single-trial firing rates at each contrast level.

Neurons with c50 estimates exceeding 95% contrast or Rmax below 1 spike/sec in more than half of bootstrap iterations for either condition (i.e. nicotine or saline) were excluded (4/121). We then used Wilcoxon rank-sum tests to compare the 1000 saline and nicotine bootstrap values of Rmax and c50 for each neuron, with p-values corrected across all units (Benjamini-Hochberg, FDR alpha = 0.05). Direction was assigned from the ratio of bootstrap medians. Model-predicted Rmax and c50 were obtained by fitting a Naka-Rushton function to the predicted response magnitudes.

#### Non-parametric analysis of contrast responses

We obtained an approximate ‘area under the curve’ measure, which we call the normalized response area (RA). To obtain this measure, we calculated the average firing rate response to each stimulus contrast, subtracted from this the average firing rate to a blank screen, and then normalized to the firing rate response for the highest stimulus contrast (96%) under the control condition relevant for the comparison (no drug baseline or pressure-matched saline). We then summed these normalized, ‘blank-subtracted’ spike rates across all contrasts, and expressed them as a ratio of the experimental to control condition (e.g., nicotine divided by saline; N:S RA, Disney et al., 2007). Both parametric (t-test) and non-parametric (Wilcoxon signed-rank) statistics were calculated and exact p-values reported.

#### Classification of units as ‘enhanced’ and ‘suppressed’

For each unit, a parametric bootstrap (1000 iterations) drew a firing rate for each contrast from a Gaussian defined by the statistics of the contrast responses for that individual neuron in the nicotine trials. The normalized N:S RA was computed as above for each iteration; the unit was classified ‘enhanced’ if the 95% CI excluded N:S RA = 1 from above, ‘suppressed’ if from below, ‘unchanged’ otherwise. The resulting counts (of the number of enhanced or suppressed units) were tested against a per-unit false-positive floor determined by splitting each unit’s saline trials into random halves and applying the same bootstrap and CI to the half-to-half ratio, over 200 re-randomizations. This yielded a 5.26% false positive rate against the nominal 5% (α = 0.05). For layer 4C, where nicotinic receptors (cation channels) are expressed on glutamatergic axons, we applied an *a priori* one direction binomial test (α = 0.05). Outside layer 4C, where there is ∼0% high-affinity nicotinic receptor expression and no response to nicotine (Disney et al., 2007), the expected direction of effects is not defined; thus a two-direction test was applied (α = 0.025 on each tail).

### Normalization Model of Attention

We used both publicly available scripts (https://snl.salk.edu/∼reynolds/Normalization_Model_of_Attention/) and our own custom MATLAB code for this analysis.

#### Using port tuning as priors on Ax, Aθ

The fitted nicotine gain field center was constrained by independent measurements at the fluid port. The spatial center (Ax) was initialized from the RF visual field position for channels 16 and 17, which flanked the fluid port. The orientation center (Aθ) was similarly initialized from orientation tuning at these electrodes 15-18.

As a post-hoc plausibility check on the fitted gain field width (AθWidth), we conducted a port tuning survey across channels 14–19. Each channel’s tuning estimate was assigned both a confidence weight (high = 1.0 for well-isolated units with clear tuning; low = 0.5 for noisy, weakly tuned, or multi-unit recordings; none = 0 when no responsive units were present) and a distance weight from the port (1.0 for channels 16–17 immediately flanking the port, and 1/d for more distant channels). The circular variance of the weighted orientation preferences provided an empirical estimate of orientation preference peak scatter at the delivery site.

#### Fitting the model

We fit the normalization model to the observed nicotine/saline response ratios using a hierarchical optimization procedure. Parameters were divided into two classes: per-session parameters describing the nicotine gain field for each recording day (peak gain, Apeak; orientation center, Aθ; orientation tuning width, AθWidth; and spatial center, Ax), and global parameters shared across all sessions (saline spatial and orientation gain field width: salAxWidth and salAθWidth, respectively; and excitatory and inhibitory kernels for model unit orientation tuning: EθWidth and IθWidth, respectively). The model was thus fit with 4 free parameters per session (Apeak, Aθ, AθWidth, Ax) and 4 shared parameters across sessions (salAxWidth, salAθWidth, EθWidth, IθWidth), totalling 36 free parameters for 121 observations. The nicotine spatial width (nicAxWidth) was fixed at 2.4°; the convention in the original model publication for a regime in which the number of neurons exposed to the stimulus field is much larger than the number subject to the attention field was attention field = 0.6*stimulus. We are firmly in such a regime in this experiment; in our model space, stimuli are 4°, yielding a field size of 2.4° per convention. A sensitivity analysis sweeping nicAxWidth from 0.5° to 8° (see Results) confirmed that sign accuracy was largely insensitive to precise values across this range.

Per-day drug parameters and shared global parameters were jointly estimated by alternating block-coordinate optimization. The objective was a weighted sum of softplus sign-classification losses on two predicted readouts — the nicotine:saline response ratio at 96% contrast and the nicotine:saline ratio of normalized RAs—with an L2 penalty regularizing fitted Ax and Aθ values toward port-tuning priors (described above).

Each block was solved by bounded sequential quadratic programming (MATLAB fmincon, SQP algorithm), with a two-stage warm start within each per-day block: stage 1 held Apeak for nicotine fixed at the saline value (4) and fit only AθWidth, Ax, and Aθ; stage 2 released Apeak for nicotine with bounds [1, 8].

For each recording day, each neuron’s position in model space was determined by a) projecting its receptive field location onto a ‘stimulus axis’ (a vector from central fixation through the center of the visual stimulus position), and b) computing its preferred orientation relative to the stimulus orientation. The model predicted nicotine:saline response ratios (on peak contrast and RA) for each neuron by computing the full normalization equation under both nicotine and saline gain field configurations across all tested contrasts. The loss summed two softplus sign-classification penalties—one on the predicted N:S ratio at 96% contrast, one on the N:S ratio of normalized RAs—using a slope of k=10 around classification thresholds of 0.99 and 1.01. Each neuron’s observed response was first labelled as increased (N:S > 1.01), decreased (< 0.99), or unchanged; the softplus penalty pushed each predicted ratio into the correct band. Units whose observed ratio fell within a narrow crossover band [0.9, 1.1] were upweighted by a factor scaling linearly with their distance from the fitted gain-field center (capped at 2×), so that units near the gain field boundary contributed more to constraining its spatial and orientation extent. Optimization proceeded in three stages: first, independent per-session fits were performed with global parameters fixed at initial estimates; second, global and per-session parameters were alternately optimized, iterating until convergence; and third, a final per-session refit was performed with the converged global parameters held fixed.

#### Direct versus cascade models for infragranular units

To test whether infragranular responses could be better explained by a model that reflects the canonical feedforward circuit from supragranular to infragranular layers, we compared two approaches for predicting infragranular effects. In the ‘direct’ model, infragranular units were treated identically to supragranular units: each unit’s predicted response is determined only by its position relative to the nicotine gain field. In the ‘cascade’ model, infragranular neurons receive as excitatory drive the output of the supragranular population after normalization. In this cascade model, the nicotine effect on infragranular units is inherited indirectly: the gain field reshapes the supragranular population response surface, and that reshaped output serves as the drive to a second normalization stage with no direct gain field of its own. The two models (direct, cascade) were then blended by a mixing parameter alpha, where alpha = 1 recovers the direct model and alpha = 0 represents the pure cascade (no direct gain field effect). For the cascade model, the gain field was fit to the supragranular units to which it was applied: at each point of a 2D grid over (EθWidth, IθWidth), per-day parameters were fitted to supragranular data only; infragranular sign accuracy was then evaluated as a function of α ∈ {0, 0.1, …, 1.0} at each grid point. Infragranular predictions were therefore entirely out of sample with respect to the parameters being used. We report results at the grid point that maximized supragranular accuracy.

#### Evaluating model performance

To assess whether the normalization model captures genuine structure in the data (rather than achieving accuracy by overfitting to the base rates of enhancement and suppression) we performed a label-shuffle permutation test. Model predictions were computed once, using the fitted per-day and global parameters. For each of 10,000 permutations, the observed effect labels (increased, decreased, or unchanged) were randomly shuffled across units independently within each recording day, preserving the per-day proportions of each class. Sign accuracy was then computed between the fixed predictions and the shuffled labels.

To assess the contribution of orientation tuning to model performance, we compared the full model against a position-only variant in which the orientation bandwidth (AθWidth) of the nicotine gain field was set to 180° (uniform across all orientations). All other parameters were held at their fitted values. Sign accuracy and Spearman correlations were computed for both models using the same classification thresholds and compared to the shuffled-label null distribution.

## Results

Nicotinic acetylcholine receptors (nAChRs) are expressed at thalamocortical synapses across species and sensory systems, and their activation strongly enhances gain (Prusky et al., 1987; Gil et al., 1997; Kimura et al., 1999; Hsieh et al., 2000; Disney et al., 2007; Kruglikov and Rudy, 2008). However, most of these studies were undertaken *in vitro*; the layer 4C gain enhancement under delivery of nicotine, that we characterized *in vivo* under anesthesia (Disney et al., 2007), has not yet been demonstrated in the awake animal. We thus first assayed the local (within-layer) effect of activating nAChRs in layer 4C of V1 in awake and fixating rhesus monkeys, using neural contrast responses to Gabor stimuli as a measure of gain (**Fig. 1**).

We used 32-channel electrode arrays (Plexon, S-Probe) equipped with a fluid capillary (**Fig. 1D**) to deliver nicotine and simultaneously record from the entire cortical depth of V1 across 9 sessions (4 from Monkey D, male; 5 from Monkey E, female). We positioned the fluid capillary exit (‘fluid port’) near layer 4C based on a current source density (CSD) analysis of visually-evoked local field potentials (**Fig. 1D**).

### The local effect of nicotine in layer 4C

After excluding recording sites that were visually unresponsive, poorly tuned, and/or unstable under pressure ejection (see Methods), we had 160 unit recordings (Monkey D: 75; Monkey E: 85) available for our main analysis of nicotine effects, and a further 28 recordings from a stationarity control study (Monkey E). Of these mostly multi-units (MU), 77 were in the supragranular layers (2-4B), 39 in the granular layer (4C), and 44 in the infragranular layers (5, 6).

We saw an increase in layer 4C spiking activity when we delivered nicotine (**Fig. 2A,B**), similar in magnitude and form to the gain enhancement observed in layer 4C of macaque V1 under anesthesia (Disney et al., 2007). Nicotinic receptors are ligand-gated cation channels and are expressed on (glutamatergic) thalamocortical axons presynaptic to excitatory layer 4C neurons (Disney et al., 2007); increased firing is the expected direct effect. We also observed decreases in firing. Attributing any of these bidirectional local (layer 4C) changes to a direct effect of nicotine requires demonstrating 1) recording stationarity and 2) a lack of gain changes during delivery of the diluent (saline).

**Figure 2.**
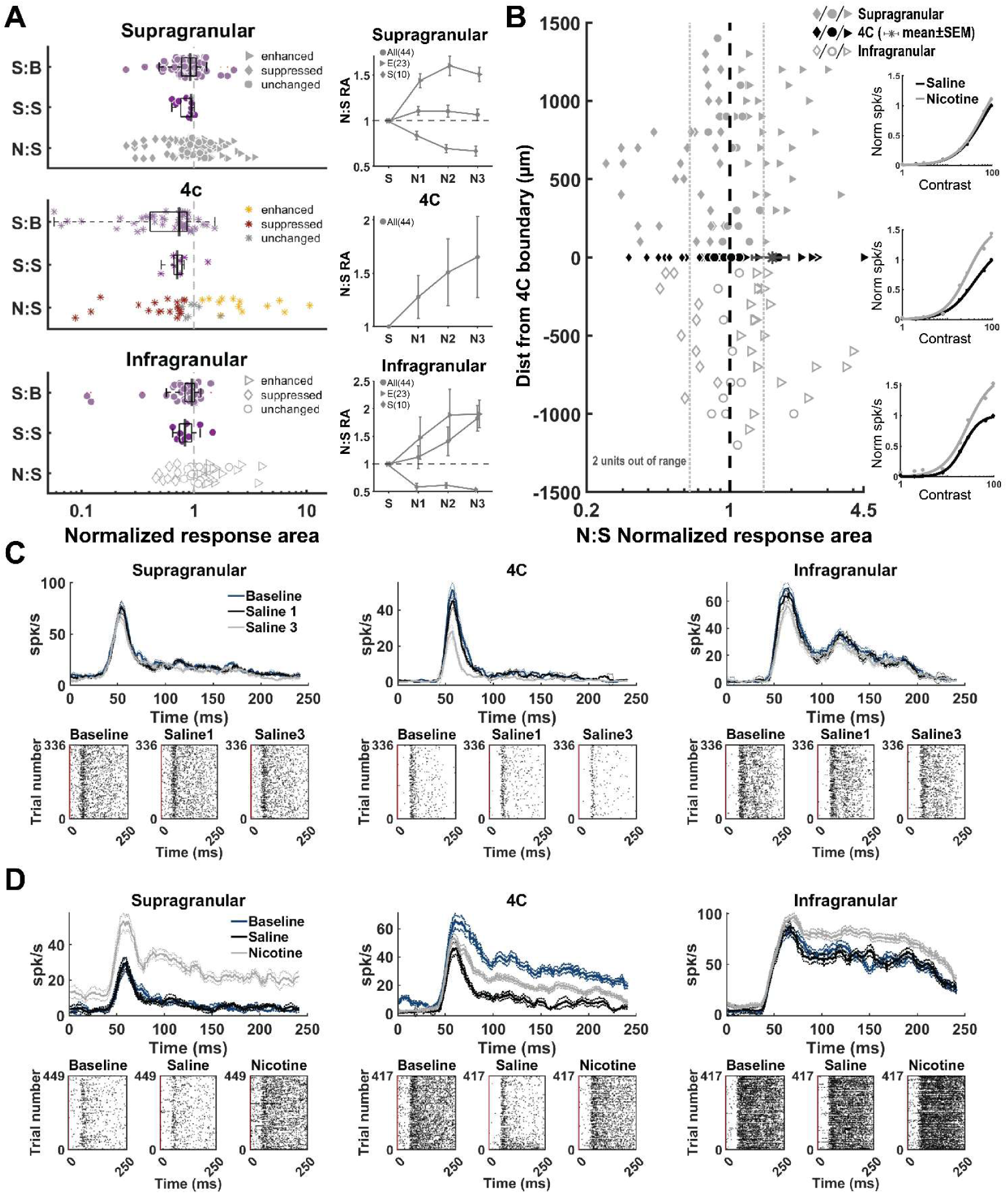
Nicotine enhances gain in layer 4C, and the effect propagates. **(A)** RA ratio by laminar category and condition. Boxplots: median and IQR. Light purple: saline to (no fluid ejection) baseline, S:B; dark purple: saline stationarity study, first to last saline block, S:S; Gray (red/gray/gold for 4C): nicotine to saline, N:S, with RA bootstrap categorization indicated by shape (color for 4C): enhanced (triangle/gold), no effect (circle/gray), and suppressed (diamond/red). Insets: ‘Dose’ response curves; population (direct) nicotine effect in 4C; broken out by enhanced (triangles), suppressed (diamonds) and all (circles) in the extragranular layers. **(B)** Nicotine to saline (N:S) response area (RA) as a function of distance from the layer 4C boundary. The specific electrode contacts assigned to layer 4C (∼6, determined by CSD) varied across sessions; depth in panel B is therefore expressed relative to the CSD-defined 4C boundaries (i.e., all within 4C contacts collapsed to depth = 0). Dark gray asterisk: layer 4C mean ± SEM (n=39). Extragranular layer symbols as in (A). Black filled symbols: individual 4C units. Dashed line: RA = 1, i.e., no difference in normalized, blank-subtracted, total spike count between the conditions. Dotted lines indicate the magnitude range observed in the stationarity control (S:S RA) study. Main plot n=160 units (nicotine days only), 2 animals. Insets: population contrast response function fits per layer (gray: nicotine; black: saline). **(C)** Example recordings showing stationarity under repeated saline delivery. Top: spike density functions (sliding window average, ± SEM) for no fluid delivery baseline (dark blue), first saline (black), and last saline (light gray) block. Bottom: Raster plots aligned to stimulus onset (t=0; red line). **(D)** Example recordings showing nicotine effects. Conventions as in C; light gray is the pressure-matched nicotine condition.

### Recordings are stationary during saline delivery

Delivering fluids by pressure ejection can cause a loss of unit isolation hundreds of microns away (Dagdeviren et al., 2018; Garwood et al., 2023). Each recording day, we started by measuring neural contrast responses over a series of saline controls with increasing ejection pressure (S:B RA, **Fig. 2A**). Detected spikes decreased over these trials. The magnitude of this apparent ‘suppression’ (loss of unit isolation) was correlated with distance from the port (Spearman’s ρ; p = 0.03) in the extragranular (i.e., supragranular and infragranular) layers, and was uncorrelated distance from the port within 4C (p = 0.25). The nicotine effect we report below shows the opposite pattern (see below). There is thus a dissociation between the mechanical artifact and drug effect. We lost all but 3 well-isolated units during these controls; using MU activity to assay pharmacological manipulations that have bi-directional effects (as ours do, see below) via sparsely expressed receptors is conservative: multi-unit activity averages over units of mixed response types and thus attenuates net effects; a change visible in the population measure likely underestimates single-unit effects.

Importantly, we found that this loss of unit isolation was a one-off displacement of the tissue with respect to the probe. Once the tissue settled in the new position, and with carefully optimized delivery parameters (see Methods), recording stationarity was excellent after the first few saline ‘pulses’ (**Fig. 2**; note range of S:S values in 2A). For each day, we capped the backing pressure for nicotine at the highest pressure that yielded stable recordings with saline and set spike sorting parameters from this stable saline condition, which serves as the control in all analyses below. On a dedicated, saline-only ‘stationarity control’ day (in which we conducted a full experiment, but just delivered saline all day, no nicotine), this stability lasted the full ∼2-hour recording (28 units; S:S RA **Fig. 2A; Supplemental Fig. S1**).

### Nicotine selectively alters visual responses

‘Spontaneous’ activity in layer 4C (mean response to blank screen trials interleaved amongst the sequence of Gabor stimuli) was unchanged by nicotine: saline average spikes/sec = 6.93 (sd = 9.06); nicotine = 7.21 (sd = 9.64; paired t-test: t(38) = -0.267, p = 0.7908). For visual responses, a contrast response function (CRF; **Fig. 2B**, insets) quantifies the relationship between stimulus intensity and spiking response. As a non-parametric measure of gain, we calculated a ‘ratio of normalized response areas’ (RA) under the CRF: for each contrast, under two conditions to be compared, we calculated the average firing rate, subtracted the blank-screen response, normalized to the 96% contrast response in the control (baseline or saline) condition, summed across contrasts, and expressed the result as a ratio (e.g., nicotine/saline; N:S or first/last saline block S:S; Disney et al., 2007).

On the background described above—i.e., weak, stable loss of single unit isolation—the average ratio of the pressure-matched nicotine and saline conditions (N:S RA) in layer 4C increased across the population on (N:S RA = 1.610; SEM ± 0.33; asterisks, **Fig. 2A,B**).

Most neurons in layer 4C are not post-synaptic to an axon expressing a nicotinic receptor (Disney et al., 2007); accordingly the <u>median</u> N:S RA in layer 4C across the nicotine recording days was identical to the median S:S RA in the stationarity control study (both median = 0.82), reflecting a dominant, stable fluid ejection artifact. To identify effects on the subpopulation of neurons post-synaptic to a nicotinic receptor-bearing axon, we classify 4C units as individually ‘enhanced’ or ‘suppressed’ using a bootstrap analysis of the N:S RA: if the 95% confidence interval excluded 1, the unit was classified as enhanced (>1) or suppressed (<1). Given that the median N:S RA is <1, this is conservative with respect to identifying the population of interest (i.e., neurons directly responsive to nicotine). By this metric, 13 (of 39) layer 4C units were enhanced, versus the one unit expected under an empirically calibrated false-positive floor (one-sided binomial test, p = 6.6 × 10⁻¹²). For the 19 layer 4C units that were classified ‘suppressed’ by this metric, we cannot separate mechanical artifact from a circuit effect downstream of the direct effect of nicotine. Effect magnitude within layer 4C was correlated with distance from the drug port (Spearman’s ρ = −0.41, p = 0.0096).

We tested up to three nicotine ‘doses’ (backing pressure levels); the population N:S RA in layer 4C increased with dose (**Fig. 2A**, middle inset). Nicotine is lipophilic and persists after ejection; N:S RA did not return to baseline over the ∼7-15 minutes of recovery possible within a recording day (recovery- saline, paired t-test: t(38) = 2.724, p = 0.01259). Consistent with a ligand-specific effect, the lack of recovery was restricted to enhanced units (recovery-saline: t(12) = 3.333, p = 0.002887); the 19 suppressed units <u>did</u> recover (t(18) = 0.845, p = 0.3912), which is not consistent with continued loss of unit isolation or injection-related tissue damage.

### Delivering nicotine to layer 4C has heterogeneous effects in other layers

The magnitude of the direct RA enhancement in layer 4C with nicotine delivery was similar to that in our prior study under anesthesia (Disney et al., 2007), so we next asked what happens to visual responses in the supragranular (2-4B) and infragranular (5,6) layers in response to this enhancement. Importantly, local nAChR expression is negligible outside layer 4C in V1, and we have shown in our earlier study that units in these layers generally do not respond to nicotine delivered locally (Disney et al., 2007). These neurons can, however, receive the modified layer 4C activity as feedforward drive.

As in layer 4C, nicotine did not change spontaneous activity for supragranular units (saline = 9.74 spikes/sec, sd = 10.64; nicotine = 10.82, sd = 12.66; t(76) = -1.062, p = 0.2916). However, nicotine <u>did</u> increase spontaneous activity in the infragranular layers (saline = 6.42 spikes/sec, sd = 6.36; nicotine = 9.54, sd = 9.62; t(43) = -3.103, p = 0.0034). Spontaneous activity in the infragranular layers did not recover (t(43) = -2.60, p = 0.0127), and the change was confined to enhanced units, i.e., those for which <u>blank-subtracted</u> (visually driven) activity was also increased.

The effect on visually-driven activity in the extragranular layers of the gain change in layer 4C visual responses varied enormously (N:S RA range: 0.13-3.98). By bootstrap analysis, we find that 95 of 121 neurons <u>outside layer 4C</u> show individually significant changes when nicotine is delivered hundreds of microns away, inside layer 4C (95% bootstrap CI excludes 1). We observed 53 enhanced and 42 suppressed MU, versus ∼6 expected by chance (binomial p ≈ 10⁻⁹⁸). Amongst these units, there were 30 enhanced (median N:S RA = 1.768) and 32 suppressed (median = 0.666) supragranular units, with 15 showing ‘no net effect’ (see below). In the infragranular layers, 23 units were enhanced (median N:S RA = 1.367), 10 were suppressed (median = 0.825), and 11 showed no net effect. Response magnitudes in the extragranular layers during nicotine delivery (N:S) were uncorrelated with distance from the drug delivery site (supragranular Spearman’s ρ = −0.10, p = 0.40; infragranular ρ = −0.15, p = 0.32; **Fig. 2B**). This contrasts with changes due to mechanical artifact, as assayed by the saline to baseline (S:B) RA, which were significantly correlated with distance (reported above). There is thus a dissociation between the drug effect (correlated with distance within 4C, but not outside) and the mechanical artifact (correlated with distance outside 4C, but not within).

Total volumes delivered (saline + nicotine) over the hours-long recording days were small (<<2uL) and recording sites on different days were well separated over the V1 surface, limiting effects of cumulative damage. Accordingly, N:S RA magnitude was not correlated with recording chronology (animal D: ρ = -0.09, p = 0.47; animal E: ρ = -0.20, p = 0.06) and rates of ‘suppression’ specifically did not increase over time (animal D: ρ = +0.07, p = 0.56; animal E: ρ = +0.09, p = 0.43). Given our permissive inclusion criteria and ‘best fit’ stimulus (that leads to low drive/firing rates in some neurons), we performed a sensitivity analysis on exclusion of low firing rate neurons: the per-unit bootstrap classification was rerun after excluding units at three thresholds (≥5, ≥10, ≥20 spikes/sec response to the 96% stimulus contrast with saline, excluding 10%, 19%, and 31% of units respectively). All primary population tests remained significant at all thresholds in all three laminar groupings (granular/4C, supragranular, infragranular); the 4C gain enhancement foundational to the study claims is still significant (p = 3.5×10⁻⁶) at the ≥20 Hz cutoff.

It is important to note that the ‘no net effect’ classification (26 neurons) outside layer 4C is distinct from the ‘no effect’ class in 4C: extragranular neurons are not directly affected by nicotine, so an unchanged N:S RA reflects precisely balanced enhancement and suppression (see normalization model below); no *net* effect.

Dose response curves within gain change classes were roughly monotonic (**Fig. 2A**, insets), with some ‘saturation’ (changing enhancement/suppression balance). Unit-by-unit analysis revealed 5 enhanced units (3 supragranular, 2 infragranular) with inverted U-shaped dose responses. In the analyses below, we use the same pressure-matched nicotine condition for all units; for these 5 units this is not the peak response during nicotine delivery.

### Neither stimulus features nor receptive field offset explain extragranular responses

We wondered to what extent the pattern of response changes across the population might be explained by individual units being strongly versus poorly driven by our chosen ‘best match over the array’ stimulus (see Methods). <u>Neither change magnitude nor direction</u> was correlated with the difference between a unit’s preferred orientation and the stimulus orientation (**Fig. 3A**), nor with receptive field (RF) overlap (**Fig. 3B**). Note that units with 100% RF overlap showed the full range of N:S RA magnitudes (**Fig. 3B**, inset). This result is not changed if we restrict the analysis to ‘well-driven’ units (see **Supplemental Fig. S2**). This is unsurprising because the difference between each unit’s tuning and the on-screen stimulus is held constant across the nicotine and saline conditions and so cannot, by itself, predict a difference between them.

**Figure 3.**
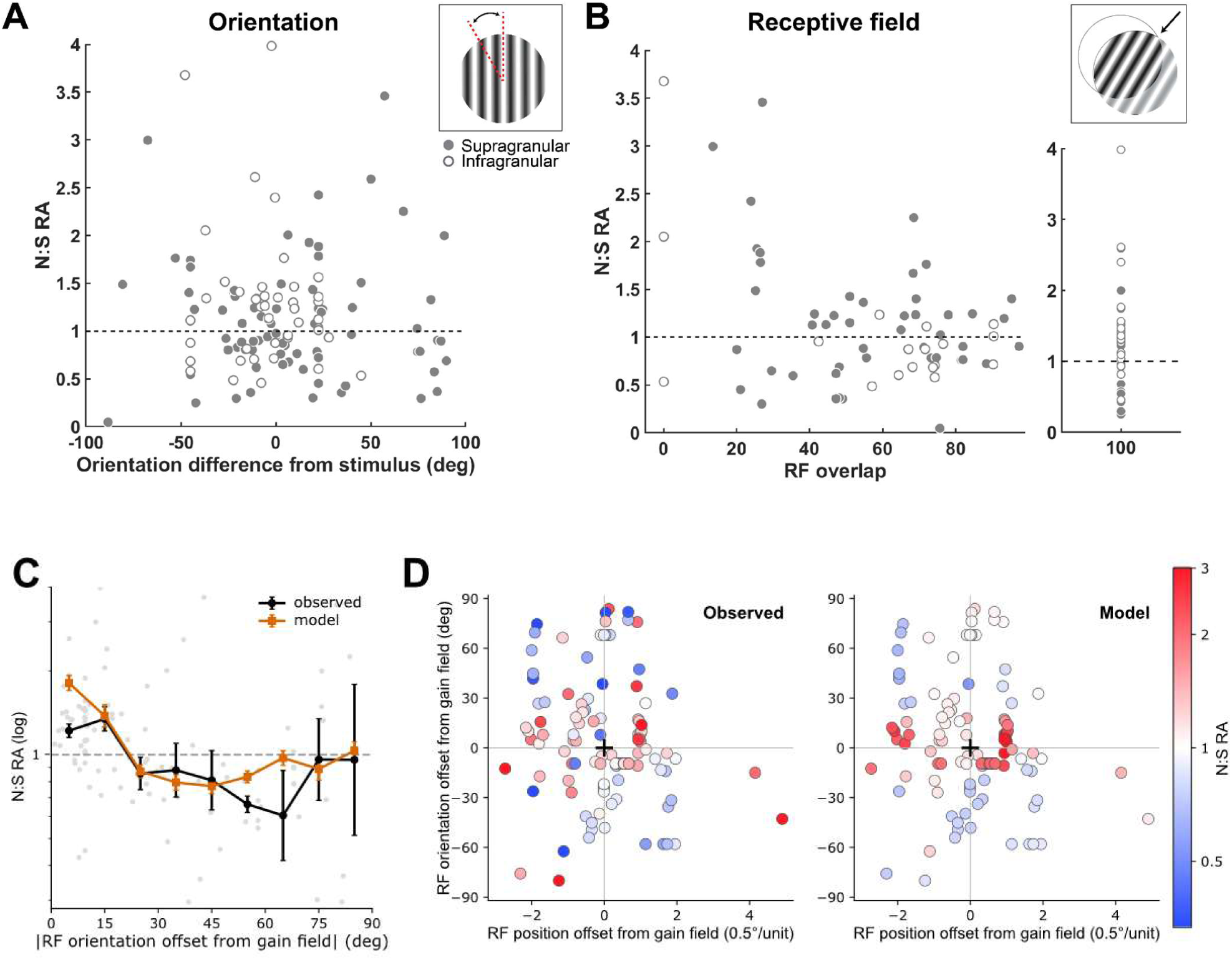
Response during nicotine delivery is not predicted by stimulus position or features, but by the offset between the gain field in layer 4C and a recorded neuron’s RF tuning. **(A)** Response area (N:S RA) vs. orientation offset between the RF and the stimulus (degrees). Filled: supragranular (n=77); open: infragranular (n=44). Dashed line: RA = 1. **(B)** Response area vs. RF overlap. Inset: units with 100% overlap. **(C)** Response area as a function of offset between a unit’s RF tuning for orientation and the orientation tuning (Aθ) of the gain field fitted to the normalization model (|RF preferred orientation − gain-field orientation|). Gray points, individual units; black, observed; orange, model-predicted; geometric mean ± SEM in 10° bins. Dashed line: RA = 1. **(D)** Each unit is plotted at its offset from the gain field with respect to RF position and preferred orientation, colored by N:S RA magnitude and direction (blue, suppressed; white, no change; red, enhanced). Left, observed; right, model-predicted; ‘+’, gain-field center. Observed enhancement (left) concentrates near the gain field in the joint position × orientation space; the model captures the same 2D structure (observed ρ = −0.30, p = 0.002; model ρ = −0.55, p < 10⁻⁹).

### Applying a normalization model of attention

V1 is widely understood to be a normalizing circuit. We hypothesized that if the layer 4C gain change is reaching the extragranular layers by propagating through that circuit (rather than by a direct effect of nicotine outside layer 4C) extragranular responses should carry a signature of having been run through that normalizing circuit. Normalization models of attention are well-suited to this test (Reynolds and Heeger, 2009; Lee and Maunsell, 2010) as they implement a multiplicative gain field applied to the stimulus, which is the effect we observed for nicotine acting directly in layer 4C.

We applied Reynolds and Heeger’s implementation of a normalization model of attention (Reynolds and Heeger, 2009), using the authors’ publicly available code (see Methods). In their model, the response, R, of a neuron with RF visual position *x* and preferred orientation θ is:

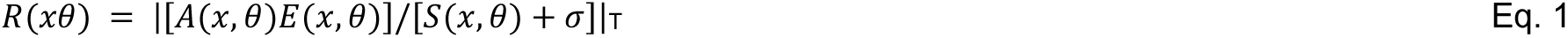

This roughly states that the response R(x,θ) is proportional to excitatory drive E(x,θ) scaled by an attention (here, nicotine) gain field A(x,θ), and divisively normalized by a pooled suppressive drive S(x,θ) plus a semisaturation constant σ. In this formulation, the normalization pool sees the product of attention and stimulus, not each independently. The multiplicative relationship between the gain field and excitatory drive has proved difficult to test causally: methods like optogenetic stimulation add stimulus-independent activity (effectively [*A*(*x*, *θ*) + *E*(*x*, *θ*)]). However, in activating receptors selectively expressed on thalamic axons, in a manner that yields a response gain increase on the visual response only, we achieve the specified multiplicative [*A*(*x*, *θ*) ∗ *E*(*x*, *θ*)] interaction.

Our model adaptation accounts for the fact that, unlike traditional attention tasks in which the stimulus field (excitatory drive) and attention field are co-located (as a result of ‘attending to’ the stimulus), our stimulus and gain fields are misaligned **(Fig. 4A**). On a given day, the population of recorded neurons vary in position and tuning offset with respect to <u>both</u> the stimulus and the gain field applied in layer 4C, and all of these misalignments differ in form across days (**Fig. 1D,E**; **Fig. 4**). Thus, to apply the model, we treated nicotine as delivering a gain field, the tuning of which was determined by the layer 4C neurons nearest the fluid port on the electrode array. This gain field therefore has its own preferred orientation, set by those layer 4C neurons’ RF; on most recording days this gain field tuning differed from the stimulus position and orientation as a result of our using the best ‘consensus’ stimulus we could find for the RFs we observed across the array. Thus, the offset between a unit’s RF and the stimulus (which does not explain the data, **Fig. 3A,B**) and the offset between that RF and gain field, differ.

**Figure 4.**
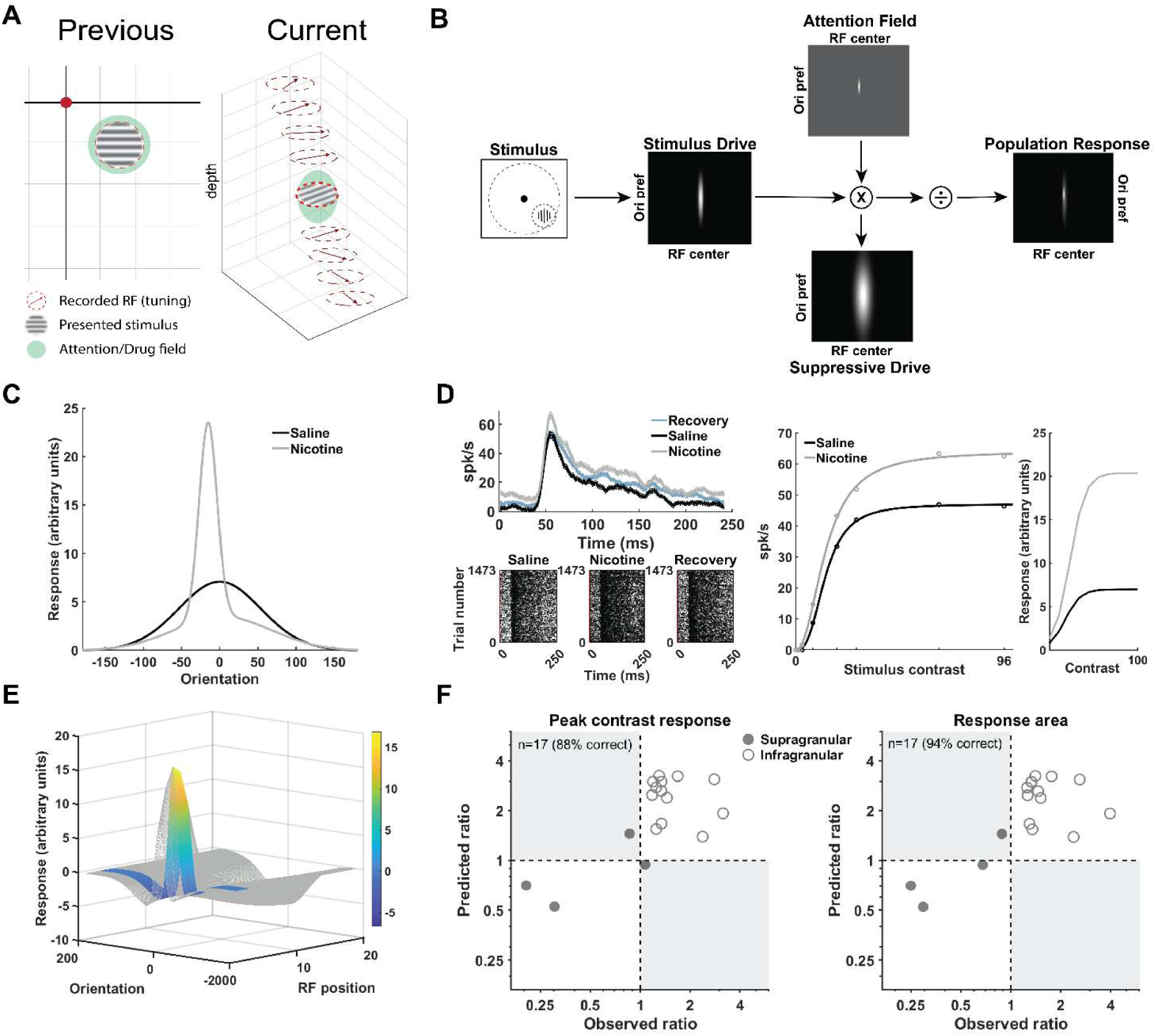
Adapted normalization model, example day (Dec 10; Monkey D). (A) Schematic comparing previous combined *in vivo* recordings (and model convention; Previous) in which RF, stimulus, and gain field are aligned with the current experiment in which they are not (Current). **(B)** Model schematic prepared using code from Reynolds and Heeger (2009) with the per-day fit parameters for Dec 10 shows a highly localized gain field (Attention Field’, top) in position (x) and orientation (θ) arising from delivery of nicotine to tissue representing a particular position in visual space and orientation preference. **(C)** Predicted gain curves at the stimulus position (x=4). **(D)** Example enhanced unit. Left: spike density functions (sliding window average ± SEM; top) and rasters (below). Center: observed contrast responses, with fits. Right: model-predicted contrast response functions. Black: saline; gray: nicotine; blue: recovery. **(E)** Full predicted gain change (nicotine minus saline) surface over orientation and receptive field position; colored region: range sampled by our array. **(F)** Predicted vs. observed ratio (N:S) for the response to 96% contrast (left; 88% correct, 15/17; Spearman’s ρ = 0.481, p = 0.0508) and RA (right; 94% correct, 16/17; RA: ρ = 0.516, p = 0.0339). Filled: supragranular; open: infragranular.

We mapped this gain field, and each unit’s RF position and tuning preference, into the model’s coordinate space (*x*,θ). RF positions were projected onto an axis running from the fixation point through the stimulus center (yielding *x* in arbitrary units ∼2 * eccentricity). Tuning preferences were expressed as deviations from the stimulus (stimulus θ = 0°).

For each recording day, we fit four parameters describing the nicotine gain field (Table 1), optimizing them to predict the classified N:S RA for each unit (ternary labels, without regard to statistical significance: enhanced [+1], N:S RA > 1.01; suppressed [-1], N:S RA < 0.99; no net effect [0], N:S RA = 1). The size of the nicotine gain field (nicAxWidth) was fixed at 2.4° per model convention (see Methods). A further four ‘global’ parameters (shared across days) were then fit to the full dataset: gain field position (salAxWidth) and tuning bandwidth (salAθWidth) <u>for saline</u>, and model neuron orientation tuning bandwidth (EθWidth, IθWidth). A final per-day refit followed with global parameters fixed.

**Table 1:** Per-day parameters.

|  | Nicotine concentration (mM) | Gain field peak (Apeak) | Gain field position (Ax, Stimx*) | Gain field tuning (Aθ; degrees) | Gain field bandwidth (AθWidth; degrees) |
| --- | --- | --- | --- | --- | --- |
| April 26 | 500 | 5.3 | 8,8 | -5 | 12 |
| Dec 10 | 200 | 7.2 | 2,4 | -16 | 12 |
| Feb 08 | 400 | 1.5 | 4,4 | 29 | 12 |
| Jan 13 | 400 | 4.0 | 4,4 | 22 | 4 |
| Jan 20 | 400 | 1.3 | 12,10 | -26 | 4 |
| Jun 11 | 400 | 1.4 | 5,6 | -148 | 12 |
| Jun 20 | 300 | 2.2 | 3,4 | 13 | 4 |
| May 16 | 250 | 8.0 | 10,10 | -55 | 12 |

\* RF positions were projected onto an axis running from fixation through the center of the stimulus position (Stimx), see Methods. Arbitrary units (AU), $\sim 2 * \text{eccentricity}$ in degrees of visual angle.

In the final fit, three days had identical, narrow, nicotine gain field tuning bandwidths (nicAθWidth = 4); three others had broader tuning (nicAθWidth = 12). The remaining two days also had broad bandwidths (nicAθWidth = 13 and 17 for Jun 11 and Feb 8, respectively). Simplifying to two values (nicAθWidth = 4 or 12) did not importantly alter prediction accuracy (defined below; 1 newly misclassified unit of 121), so the simplified solution was adopted. These bandwidth values (i.e., 4 -17 degrees) are all within the reported scatter of peak orientation tuning preference for a V1 column (Nauhaus et al., 2012; Ikezoe et al., 2013). The global parameter fit values were: saline AxWidth = 7.5; saline AθWidth = 90°; EθWidth = 50°; IθWidth = 70° (see **Supplemental Table 1** for all global values, fixed and fit).

The result is a unique model implementation for each recording day (**Fig. 4**). For a given RF position, the model predicts a mix of enhancement, suppression, and net-zero gain effects, determined by a neuron’s orientation tuning (**Fig. 3C, 4C**). Thus, the effect ternary class and magnitude are determined by the joint offset between a unit’s RF and the gain field (**Fig. 3D**). Across RF positions, the model produces a family of predicted gain curves for the nicotine and saline conditions (curves in **Fig. 4C** are for RF position: x=4, the stimulus position). The difference (nicotine minus saline) between these curves, across RF positions, defines a surface (**Fig. 4E, 3D**). Our columnar recordings sample only a subregion of this surface (colored in **Fig. 4E**). The model-predicted gain effects can then be compared to the data (**Fig. 4D,E**). On the day presented in **Figure 4** (Dec 10), depending on the measure (N:S ratio of the firing rate at 96% contrast versus N:S RA), ∼88-94% of neurons (i.e., 15/17 or 16/17) were correctly classified. Furthermore, predicted versus observed gain <u>magnitudes</u> were correlated (96% contrast: Spearman’s ρ = 0.481, p = 0.0508, n = 17; RA: ρ = 0.516, p = 0.0339, n = 17); recall that <u>only permissive ternary labels (0.99 and 1.01 cutoffs) were fit</u>.

It did not matter whether the port was actually in layer 4C; on one recording day (May 16), the fluid port was positioned in layer 4B. This fact could be diagnosed from the fits, even if we hadn’t known it from the CSD: for each day, we used the orientation tuning at electrodes adjacent to the fluid port to set priors on gain field orientation tuning, Aθ. However, nAChRs are sparsely expressed in layer 4B of macaque V1 (Disney et al., 2007); accordingly, model fitting for May 16 converged on a value for Aθ that reflected *<u>tuning at the nearest</u> <u>layer 4C sites</u>* ∼200-300 µm away from the port (electrodes 18/19; preferred θ = 103°/106°; Aθ = -55° with respect to the stimulus θ = 157.5°; Table 1), not the tuning of the units adjacent to the port (θ = 161°/151°; Aθ ≈ 0).

Over all 8 days, the model accurately predicted ternary effect class for 83% (peak contrast response; **Fig. 5A**) or 84% (N:S RA; **Fig. 5B**) of 121 units. Accuracy dropped significantly for shuffled ternary labels (peak contrast: mean = 63.2%, sd = 3.3%, p = 0.0001; N:S RA: mean = 62.7%, sd = 3.4%, p = 0.001; 10,000 iterations; **Supplemental Fig. S3**). In our shuffling approach (randomizing ternary labels), the null distribution mean (∼63%) reflects chance agreement between predicted and observed marginal class proportions (i.e., the fraction of units for each ternary class, on each day). That the observed accuracy of 83-84% significantly exceeds this base-rate expectation indicates that the model captures unit tuning structure in nicotine’s effects, not just overall prevalence of enhancement and suppression. As in the example day above (Dec 10; **Fig 4**), although the model was fit to ternary gain class only, predicted and observed gain <u>magnitudes</u> were positively correlated across the population (peak contrast: Spearman’s ρ = 0.599, p < 10⁻¹²; RA: ρ = 0.566, p < 10⁻^10^; n = 121). The concentration of the nicotine solution differed across days (Table 1); none of the fit parameters were correlated with concentration (Apeak: ρ = −0.469, p = 0.2464; AθWidth: ρ = 0.274, p = 0.5107; Ax: ρ = 0.431, p = 0.2893; Aθ: ρ = 0.216, p = 0.6107; see Discussion).

**Figure 5.**
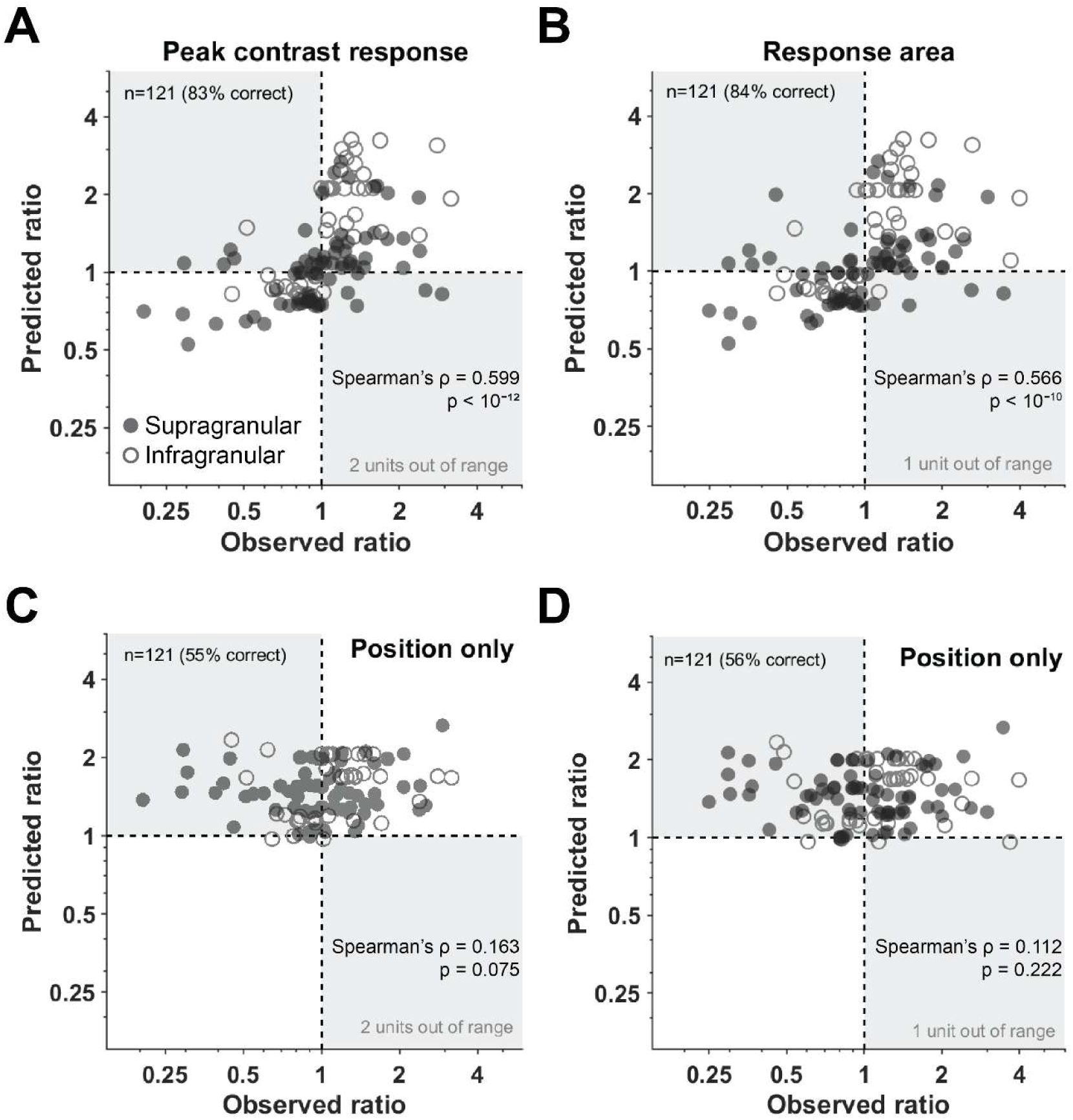
Model predicted versus observed gain changes across the population. **(A)** Observed (x-axis) versus predicted (y-axis) ratio of the response to 96% contrast. White regions: predicted and observed change classes agree; gray regions: predicted and observed classes differ. Filled circles: supragranular; open circles: infragranular. **(B)** Observed versus predicted ratio for response area (N:S RA), conventions as in A. (**C**) As in A (peak contrast prediction accuracy) for a model with position information only. (**D**) As in B (N:S RA accuracy), model with position information only.

There were 21 prediction errors (of 121 units) for peak (96%) contrast and 19 for RA. Setting a ‘within measurement noise’ tolerance at 0.8 < ratio < 1.2 (∼half the S:S RA range, **Fig. 2**), 9 of 21 peak-response errors and 4 of 19 RA errors fell within this range. Of the remaining errors, 10 were MU recordings with a poorly-defined tuning peak (i.e., secondary or broad ‘peak’). RF orientation tuning offset from the gain field is the strongest predictor of the effect of nicotine (**Fig. 3C,D**) and a sensitivity analysis (**Supplemental Fig. S4**) suggested that model performance was sensitive to gain field orientation tuning. We further evaluated a position-only variant of the model in which the gain field was untuned (AθWidth = 180°), with all other parameters unchanged. This model achieved 55.4% accuracy for the peak response (**Fig. 5C**) and 56.2% for RA (**Fig. 5D**)—well below the shuffled-label null (63.2%) and full model (82.6-84.3%). Thus, the inability to properly define a preferred orientation for these 10 poorly tuned units is expected to undermine prediction accuracy.

This leaves six well-tuned units (of 21 mis-predicted units) with substantive prediction errors for both peak contrast and RA. One was clearly lost during recording (final N:S RA 0.05). Another three had RF position and tuning preferences that placed them in an extremely low firing-rate regime for our ‘best fit’ stimulus (< 5 spikes/second). In this regime, single spikes can alter observed ternary class; all three were predicted ‘no effect’ but observed suppressed. Thus<u>, in our population of 121 neurons, there were only two well-tuned, well-driven mis-predicted units</u>: Both were in the supragranular layers. For one an enhancement was predicted and no net effect was observed (N:S peak contrast = 0.998; N:S RA = 0.933); for the other no effect was predicted and suppression (N:S peak contrast = 0.427; N:S RA = 0.4591) was observed.

### Mixed gain effect types

A parametric analysis of the relationship between stimulus contrast and firing rate can be achieved by fitting a model function to the data (insets, **Fig. 2B**; Reynolds et al., 2000; Williford and Maunsell, 2006 1167; Disney et al., 2007). We fit neural CRFs to a hyperbolic ratio function (see Methods) that yields four parameters: slope (n), asymptotic firing rate (Rmax), contrast at half maximum firing rate (c50), and offset (sFR). Changes in the Rmax and c50 parameters are often used to assess ‘response’ (Rmax) versus ‘contrast’ (c50) gain. In particular, normalization models predict that when the stimulus is larger than the gain field (as it is in our configuration), many neurons will individually show changes in Rmax and c50, but over the population, only Rmax will change.

To test this prediction against our data, we analyzed bootstrapped fits (1000 resamples) to individual neurons’ CRFs (77 supragranular and 44 infragranular) and tested for significant parameter changes using Wilcoxon rank-sum tests. After FDR correction, 110/121 units (91%) individually showed significant Rmax changes; of 119 units for which c50 was well-constrained, 80 (67%) showed significant c50 changes. The fits captured the per-unit CRFs well: the median R² for the fits were 0.89 (saline; IQR 0.81–0.94) and 0.91 (nicotine; IQR 0.80–0.96). Fits explained >90% of the variance in 46 – 51% (saline, nicotine) of units. Fits explained <50% of the variance in fewer than 8% of units (nicotine; 4% for saline). Across the population, we observed four gain change types in individual cells: response gain only (31%), contrast gain only (6%), mixed (60%), and no change (3%). As predicted by the model, despite the high prevalence of individually significant changes, the population c50 was unchanged (saline = 27.4%; nicotine = 27.6% contrast; **Fig. 6**, right; Wilcoxon signed-rank: p = 0.42; paired t test on log(nic/sal): t(118) = 0.66, p = 0.51). Also, predicted by the model is the significant population-level shift in Rmax (saline median = 40.3; nicotine = 45.4 spikes/sec; **Fig. 6**, left; Wilcoxon signed-rank (nic − sal): p = 5.3 × 10⁻⁵; paired t test on log(nic/sal): t(120) = 3.81, p = 2.2 × 10⁻⁴).

**Figure 6.**
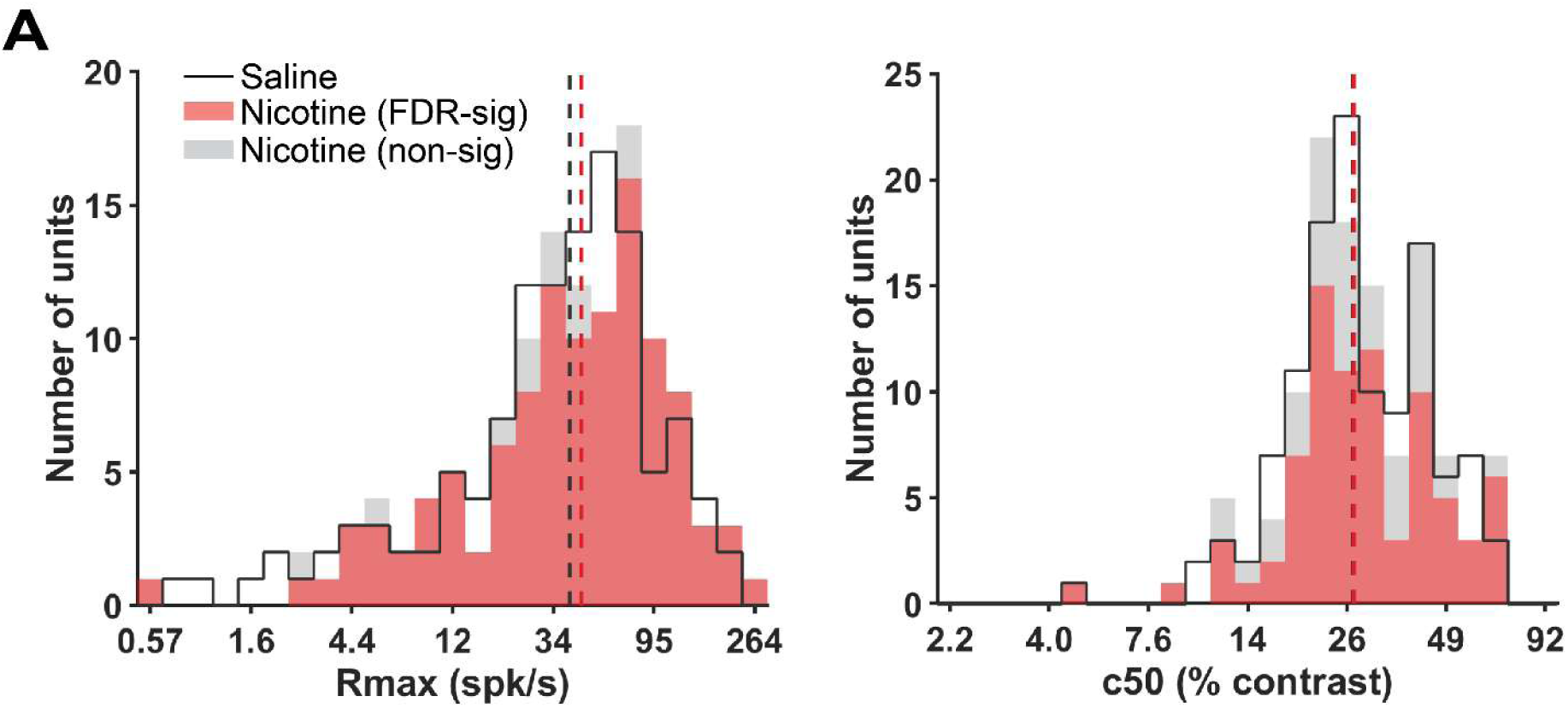
Parametric gain change analysis. Histograms of median fit values for Rmax (left) and c50 (right) in the saline (black, unfilled) condition, and for units with significant (red) and non-significant (gray) differences, collapsed over laminar categories. Black (saline) and red (nicotine) vertical dashed lines: population medians; Rmax: saline = 40.3, nicotine = 45.4; c50: saline = 27.4, nicotine = 27.6.

Our model thus far treats all extragranular units identically, essentially assuming direct input from layer 4C to all layers. We tested whether infragranular predictions would improve under a ‘cascade’ model in which supragranular unit output mediates the effect of the gain field. Refitting per-day parameters to the 77 supragranular units alone (also allowing EθWidth and IθWidth to vary; remaining global parameters fixed) yielded narrower orientation tuning than fitting to the full population (supragranular-only EθWidth = 20°, IθWidth = 50°) and modestly improved supragranular accuracy (peak contrast/N:S RA: 79%/82%, up from 78%/79%). We then generated out-of-sample predictions for infragranular units, blending direct and cascade models with a mixing parameter α (α = 1: direct model; α = 0: pure cascade). The best infragranular accuracy occurred at α = 0.9, with a prediction accuracy (93% N:S RA), identical to the direct model. However, this sign-based comparison is insensitive to the effect magnitude. The direct model (α = 1.0) systematically over-predicted the infragranular gain change (predicted median N:S RA = 1.63 vs 1.19 observed). A mixed model (α = 0.5) resolved this magnitude mismatch and improved the out-of-sample rank correlation between predicted and observed infragranular gain (Spearman ρ = 0.72, vs 0.65 for the direct model), with only a modest loss of sign prediction accuracy (drop from 93% to 91%).

### The gain effects evident in neural activity impact perceptual performance

Our gain field injection into layer 4C altered activity across the entire cortical column (**Fig. 2**) in a manner that was well predicted by a normalization model with a multiplicative gain field. We wondered whether these neural activity changes would be evident in behavior as a change in contrast perception (gain change) that is orientation-specific (normalization signature). We trained both animals on a two-alternative forced-choice contrast discrimination task designed to determine the ‘point of subjective equivalence’ (PSE) between two simultaneously presented static Gabor stimuli of matched size, orientation, and spatial frequency. The task was to saccade to the Gabor with higher apparent contrast (**Fig. 7A**). We delivered nicotine to layer 4C during task performance on 7 recording days (∼5,200 trials). The smallest number of total trials in a single session was 457, the largest 1,033; each possible contrast comparison had a minimum of 3 repeats within a single session.

**Figure 7.**
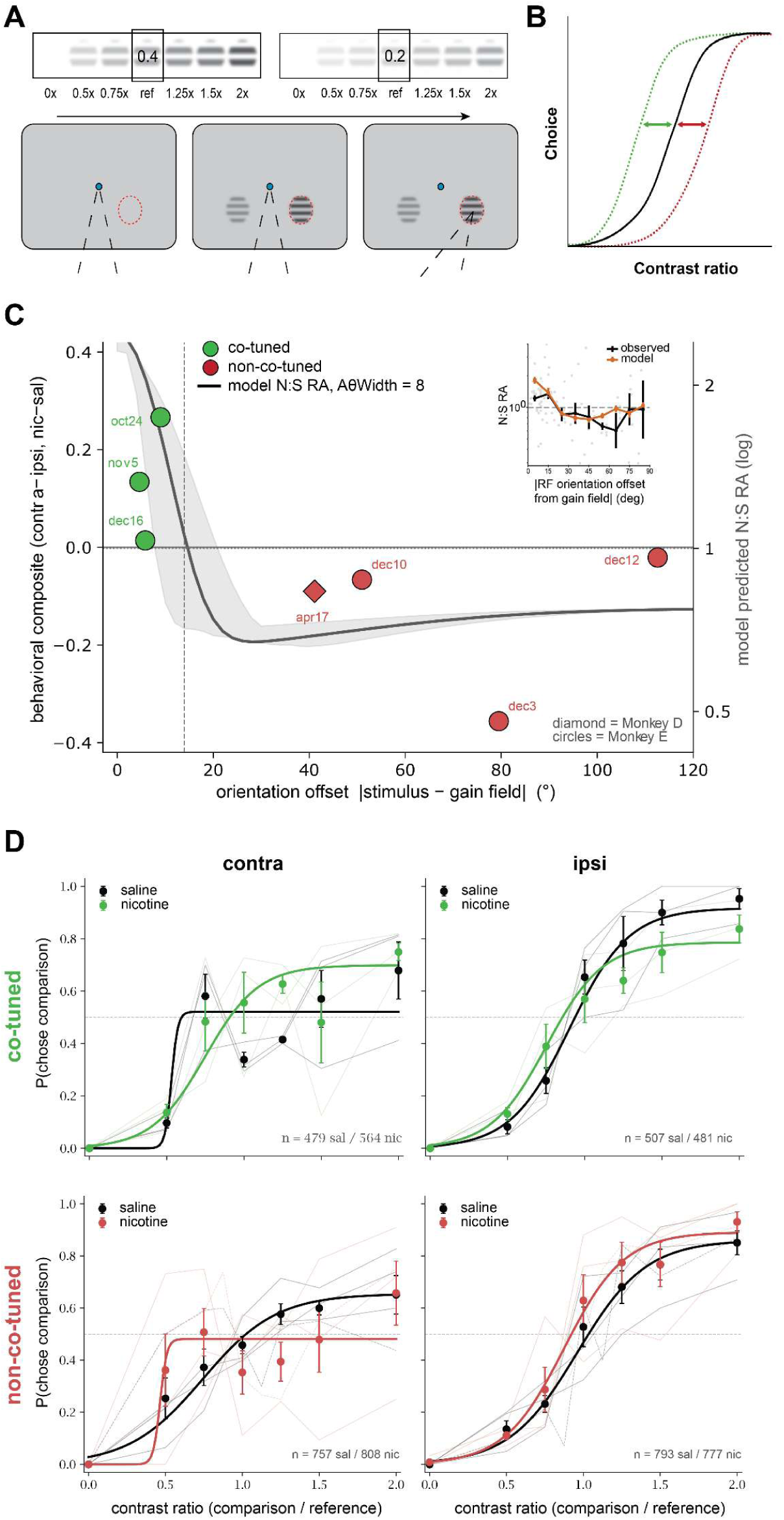
Nicotine alters perceived contrast. **(A)** Task schematic. Upper: comparison contrasts at half-steps around 20% and 40% references. Lower: trial sequence—animal fixates for a variable period of time, after which two Gabors appear (one in the gain field, red circle; one at matched eccentricity contralaterally); one is a reference contrast (0.2, 0.4), the other randomly selected from possible target contrasts. To receive a juice reward, the animal must saccade to the higher contrast stimulus. Stimuli of equal contrast are randomly rewarded. **(B)** Predicted psychometric curve shifts: leftward (green) when stimulus and gain fields are co-tuned; rightward (red) when not. **(C)** Net rightward bias shift (nicotine minus saline) vs. orientation offset. All 7 sessions shifted in the predicted direction (sign test, p = 0.016; t(6) = 2.71, p=0.035). **(D)** Choice behavior for three co-tuned (upper) and four non-co-tuned (lower) sessions. Filled circles: session mean ± SEM; open circles: individual sessions. Curve: logistic fits to pooled means. Black: saline; green/red: nicotine.

We fit the normalization model to each session’s gain field (layer 4C) tuning to predict the net expected ternary class effect as a function of stimulus orientation (gray curve, **Fig. 7C**) for these entirely out-of-sample data (behavior was collected on different days from neural data). The normalization model makes predictions for these two session types: when the gain field is co-tuned with the stimulus, nicotine delivery into V1 should *increase* apparent contrast of the stimulus in the contralateral visual hemifield, lowering the PSE and shifting the psychometric curve over contrast leftward (green, **Fig. 7B, 7D top**). When the gain field is not co-tuned, nicotine should *reduce* the apparent contrast of a stimulus in the gain field (having increased the apparent contrast for another tuning), raising the PSE, and shifting the psychometric curve rightward (red, **Fig. 7B, 7D bottom**).

To test the model predicted tuning specificity of the gain effect (**Fig. 3C, 7C** inset), on three days we selected on-screen stimuli that were within 10° of the RF tuning for the layer 4C units nearest to the fluid port (green points and curves in **Fig. 7**); on the other four days, the on-screen stimulus was chosen such that the offset between it and the tuning near the port differed substantially (red points and curves, **Fig. 7**). Six of the sessions were collected in animal E, we collected one session in animal D with tuning offset matched closely to one of the days in E (D: Apr 17; E: Dec 10), providing a strong test of effect magnitude for matched offsets across two animals. We tested the model predictions using a within-session composite measure of the net shift in choice bias under nicotine relative to saline. On each of the seven sessions, the composite score shifted in the direction predicted by the model (sign test, p = 0.016; one-sample t-test, t(6) = 2.71, p = 0.035; **Fig. 7C**). Our *in vivo* recording disrupted choice behavior in the contralateral visual hemifield, introducing a bias that was not present in either animal before or after recording (**Supplemental Fig. S5).** This bias precluded quantitative assessment of PSE shift; three-parameter logistic fits to the choice data (i.e., variable upper asymptote to accommodate side bias; **Fig. 7D**) had an asymptote < 50% choice often enough to preclude PSA calculation (see co-tuned saline and non-co-tuned nicotine conditions in the left panels of **Figure 7D**). Nonetheless, in the face of this bias against choosing the contralateral stimulus, nicotine yielded increase contralateral stimulus on co-tuned days (**Fig. 7D**, upper left), but not on non-co-tuned days (**Fig. 7D**, lower left).

## Discussion

We find that a multiplicative gain increase by nicotine — the direct effect of which is constrained by anatomy to remain within 4C — produces bidirectional changes in neuronal activity across all layers of macaque V1 that are well predicted by a normalization model with a multiplicative gain field. The same model also predicted choice on 7/7 days in an out of sample test on perceptual performance. That the responses outside layer 4C were dose responsive, were uncorrelated with distance from the site of drug delivery and bore the signature (in direction and magnitude) of having been run through a normalizing circuit supports the conclusion that these changes <u>propagated through the excitatory/inhibitory V1 circuit</u> despite having their origin in a very small number of units with modified activity in layer 4C. Thus, direct nicotinic receptor-mediated gain changes on visual processing, despite being evident in a small population of neurons, alter widespread activity. Further, this propagation does not appear to dilute or modify the tuning of the originally applied gain field.

How far does this propagation of a tuned effect go into the circuit? The model-predicted tuned effects on choice behavior suggest they may well leave V1. Numerous studies have employed pharmacological manipulations of neuromodulatory signalling in behaving monkeys, using nanoliter drug volumes, and reported changes in neural activity (Vijayraghavan et al., 2007; Wang et al., 2007; Thiele et al., 2012; Vijayraghavan et al.; Dasilva et al., 2019; Herrero and Thiele, 2021) including in V1 (Herrero et al., 2008; Seillier et al., 2017; Patel et al., 2023). In none of these studies in awake animals were behavioral effects reported.

The effect of nicotine on layer 4C responses has been reported previously (Disney et al., 2007) and is again found to be large in magnitude (mean = 1.61) but restricted to a small number of neurons (median = 0.82). It does not survive omnibus tests that treat the layer 4C population as homogenous, and thus average over it (Wilcoxon signed-rank vs 1: p = 0.71; one-sample t-test on log (N:S RA) vs 0: p = 0.87). The dominance of muscarinic over nicotinic effects in V1 as a whole has been noted often (Herrero et al., 2008; Disney et al., 2012; Soma et al., 2012; Thiele and Bellgrove, 2018; Herrero and Thiele, 2021) and the likely relevance for visual processing, attention, and perception of the very focal effects of nicotine in layer 4C in is often taken to be negligible (Hasselmo and Sarter, 2011; Vossel et al., 2014; Vijayraghavan and Everling, 2021). Our receptor-defined perturbation at the thalamocortical synapse, however, cascades through the V1 circuit and our small behavior study suggests a high upper bound on this cascade; it appears it may run far enough, without compensation, to alter perception.

The distinct patterns of expression seen for multiple modulatory receptors in layer 4, across species (Prusky et al., 1987; Rakic et al., 1988; Gil et al., 1997; Kimura et al., 1999; Hsieh et al., 2000; Watakabe et al., 2009; Zilles and Palomero-Gallagher, 2017; Disney, 2021; Rapan et al., 2022) suggests that evolution may also have ‘discovered’ this as a key site at which the brain can modify its own activity. While there are other pathways from retina to cortex, the one that runs through layer 4 (and then on through other trans-thalamic loops to other layers in higher cortices) is important for image formation. We do not know whether thalamocortical synapses are a *uniquely* powerful point for modulation; we do show a perhaps unexpectedly strong effect, and thus the value of following modulatory anatomy to test modulatory function.

Of course, manipulations that impact many neurons, such as systemic delivery (Li et al., 1999) and larger volume micro-infusions (Sawaguchi and Goldman-Rakic, 1991; Li and Mei, 1994; Noudoost and Moore, 2011a) modify behavior; the contrast between these large scale effects and those in our study is emphasized when we consider that V1 is the largest cortical area in a macaque. Our injections would have spread over a region (Martin, 1991; Tehovnik and Sommer, 1997) smaller than an orientation hypercolumn (Hubel and Wiesel, 1977; Horton and Adams, 2005).

Systemic modification of modulatory function— by experimental manipulation, drug side effects, or endogenous pathology—disrupts ‘higher level’ cognitive functions. Root causes for this are often sought in the circuits of ‘higher level’ cortex (Goldman-Rakic et al., 1990; Paspalas and Goldman-Rakic, 2005; Wang et al., 2007; Yang et al., 2013), using visual paradigms, and analyses that assume recorded neurons ‘see’ a veridical representation of on-screen stimuli. Our findings suggest that, in these studies, the visual scene that association areas ‘see’ might have been importantly modified at early stages of processing which could itself alter behavior, regardless of function or dysfunction of deeper cortical circuits. It is frequently acknowledged that systemic delivery yields organismal-level changes, but this has not yet led to systematic study of the ways modulatory systems alter the data upon which cognition and behavior depend; such studies are needed.

The capacity for nicotine, specifically, to boost local thalamocortical gain is conserved across species and sensory systems (Prusky et al., 1987; Rakic et al., 1988; Gil et al., 1997; Kimura et al., 1999; Hsieh et al., 2000; Watakabe et al., 2009; Disney, 2021); we suspect the capacity to modify sensation is also conserved, but in service of what? The bidirectional effects we observed in the supra- and infragranular layers suggest that the endogenous ligand (ACh) is not a simple ‘volume control knob’; it more likely supports complex, circuit-dependent computations, including a gain field that can be modelled by normalization models of attention (Reynolds and Heeger, 2009; Lee and Maunsell, 2010).

Normalization models of attention assert that the relationship between the attention (*A*(*x*, *θ*) in Eq. 1) and stimulus (*E*(*x*, *θ*)) fields is multiplicative. Prior studies have examined the dependence of normalization on the stimulus–receptive field relationship (Heeger, 1992; Carandini and Heeger, 2011), and an optogenetic study in macaque V1 showed that intensity-dependent normalization persists under artificial stimulation (Nassi et al., 2015). However, optogenetic (like electrical) stimulation activates neurons regardless of visual response properties (‘hijacking’ the circuit; (Griffin et al., 2011; Cheney et al., 2013), imposing an additive increase in firing rate, effectively: *A*(*x*, *θ*) + *E*(*x*, *θ*). Sanzeni et al. (2023) made this distinction in re-analyzing a V1 optogenetic dataset that was well fit on a cell-by-cell basis by the normalization model. They showed that under these conditions, the population firing rate distribution is not significantly shifted by optogenetic stimulation; instead, rates are ‘reshuffled’. Thus, despite engaging normalization at the single-neuron level, optogenetic perturbation did not implement the circuit-level gain field that normalization models of attention assume. By contrast, in applying a multiplicative response gain through endogenous receptor machinery at the thalamocortical synapse, nicotine yielded the model-specified *A*(*x*, *θ*)\**E*(*x*, *θ*). The heterogeneous responses we observed in extragranular layers are thus the predicted consequence of a focal, multiplicative gain field operating within a normalizing circuit. That our effects were well captured by a model of (endogenous) attention suggests that our perturbation engaged the circuit’s regular computational architecture; neuromodulation can reveal underlying circuit computations and perhaps also the underlying biology that supports them.

In our analysis, fitted values of AθWidth clustered bimodally with no relationship to nicotine concentration, perhaps reflecting local orientation map architecture. Narrow tuning bandwidths should be sufficient to account for a gain field applied within an iso-orientation domain; broader bandwidths might be required to account for gain applied near a pinwheel center. A future study incorporating imaging could test this hypothesis. None of the fitted parameters were correlated with nicotine concentration, consistent with gain field effects being determined by circuit features such as the orientation tuning map; we expect that the action of ACh would be similarly constrained by the receiving circuit.

We did not improve prediction *accuracy* for infragranular units with a ‘cascade’ model of V1, but when response *magnitude* was taken into account, a model that allowed for both direct and indirect (i.e., unnormalized and normalized) paths to the infragranular layers was supported. Other studies using columnar recordings with attentional modulation also support different gain effects for supragranular and infragranular layers (Nandy et al., 2017; Ferro et al., 2021). Having fit 36 parameters to 121 observations means overfitting is a concern, but our effective flexibility is lower than it sounds: 1) the model was fit to ternary class labels; 2) gain field position and tuning were constrained by independent measurements; 3) four global parameters had to explain structure across 121 units; and 4) day-specific parameters could not compensate for poor fits to other days. That observed gain magnitudes (not used in fitting) were correlated with predicted magnitudes (**Fig. 5**) suggests that the model captured real structure, as does the predicted orientation-dependent of the effect we observed on behavior, which represents an entirely out-of-sample test (different days).

Our interpretation has focused on high-affinity nAChRs; expression data for low-affinity (homomeric, α7 subunit) nAChRs is not available for macaque V1. However, radioligand binding studies indicate that low-affinity nAChRs are also selectively expressed in layer 4C (Han et al., 2003); they may well contribute to the gain effects we observed. Interestingly, unlike in macaques (or cats, or rats), in mice the nicotinic receptor expressed at the thalamocortical synapse is the low-affinity subtype (Gil et al., 1997).

ACh itself is hypothesized to support attention; could cholinergic gain control of thalamocortical transmission be ‘the’ mechanism of attention? Strictly speaking, almost certainly not: Although the capacity of the cholinergic system to specifically address such a precise volume (∼1mm) in cortex has not been determined, it would require some exotic (although biologically feasible; (Coppola et al., 2016) circuitry to do so. Herrero et al. (2008) showed that muscarinic antagonists reduced attentional modulation of V1 responses, suggesting other ACh receptors are also expressed in V1 in a way that allows them to engage endogenous machinery that supports attention. Importantly: In our study, neurons outside layer 4C did not ‘know’ we used nicotine; they just saw the gain field change. From that standpoint, we show that if you want to achieve tuned contrast enhancement of a visual stimulus (certainly one of the things visual attention does; (Reynolds and Chelazzi, 2004; Ling and Carrasco, 2006; Williford and Maunsell, 2006), gain control at the thalamocortical synapse could do the job.

That these local changes appear to impact perceptual choice implies that even modest changes to endogenous ligand levels, receptor density, or the number and morphology of thalamorecipient neurons could importantly reshape information processing. Such changes are well documented in neuropsychiatric disease: in schizophrenia, for example, alterations in cholinergic receptor expression, reduced basal forebrain volumes, and reductions in neuronal number in V1 have all been described (Butler et al., 2005; Dorph-Petersen et al., 2007; D’Souza et al., 2012; Gibbons et al., 2013; Avram et al., 2021). Our data predict that these changes, occurring even at small scales within a primary sensory area, could importantly alter the gain landscape for sensory data, and go on to bias what downstream circuits receive. The implication is not that disorders like schizophrenia are reducible to altered sensation, but that understanding neuromodulatory control of cognition, emotion, and behavior—in health and disease—likely requires that we properly account for changes taking place at the earliest stages of processing.

## Acknowledgements

This work was supported by NIH grants R01EY029663 to AAD and F32EY034772 to VCG. The authors gratefully acknowledge the assistance of L. Beaver, J. Amodeo, T. Nguyen, K. Coates and A. Brigande and the veterinary and husbandry staff of the Duke Primate Vivarium.

## Author Contributions

AAD designed the experiments, developed the adaptation of the normalization model, and assisted with analysis. VCG executed the experiments and analyzed the data. AAD and VCG co-wrote the manuscript.

## Conflict of interest statement

The authors declare no conflicts of interest.

## Notes

### Competing Interest Statement

The authors have declared no competing interest.

### Summary of Updates

additional analyses of model predictions, increased focus on phsyiology.

