## Supplemental Figures and Tables for "Nicotine causes highly localized and tuned changes in thalamocortical gain that cascade strongly and selectively through the cortical circuit"

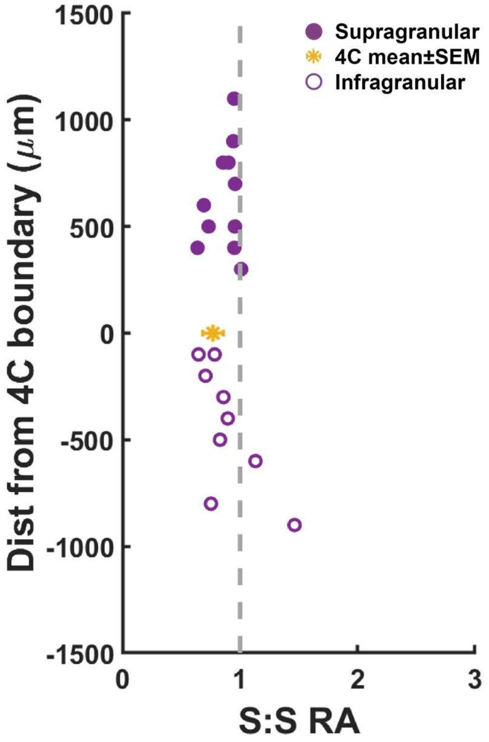


**Supplemental Figure S1. Saline stationarity control across cortical depth.** Saline was delivered in a series of trials, matching pressure doses to nicotine delivery experiments. Responses were normalized to the spike rate in response to a 96% contrast stimulus in the initial saline condition, summed, and a ratio calculated between the first and last saline conditions (S:S) to yield a response area (RA). The S:S RA (data from Fig. 2B, dark purple) is plotted against distance from the 4C boundary (4C mean± SEM in yellow) for supragranular (filled circles) and infragranular (open circles) layers. Dashed line: RA = 1, indicating no difference across saline conditions.

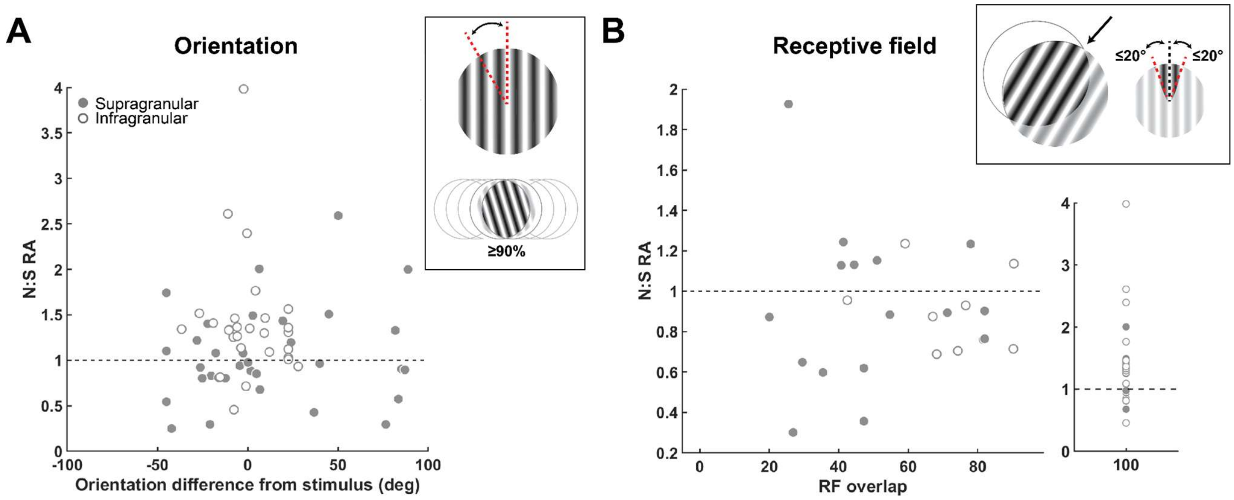


**Supplemental Figure S2. Gain change direction and magnitude are not explained by differences in tuning, even when restricted to well-tuned, strongly driven units. (A)** Normalized response area (RA) as a function of the difference between a recording site's preferred orientation and the orientation of the presented stimulus, restricted to units with at least 90% receptive field overlap. Dashed line: RA = 1, or no net nicotine effect. **(B)** RA as a function of receptive field overlap, restricted to units with a preferred orientation within 20° of the presented stimulus. Inset shows the subset of units with 100% overlap.

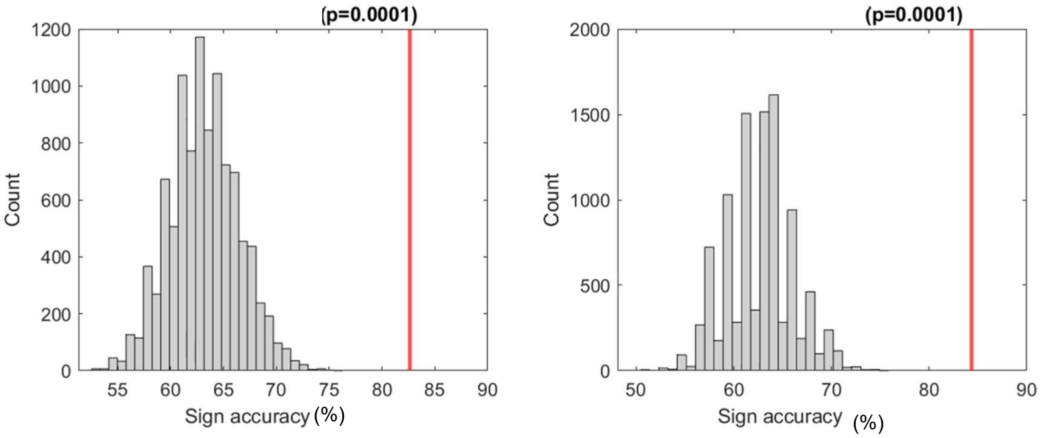


**Supplemental Figure S3. Randomization test: label shuffle within recording day.** Null distributions of sign accuracy obtained by randomly permuting the observed ternary effect labels (<0.99 → [-1]; > 1.01 → [+1], exactly 1 → [0]) across units within each recording day (10,000 permutations), while holding model parameters fixed. Gray histograms show the null distribution of sign prediction accuracy for the response magnitude ratio for the response to a 96% contrast stimulus (left) and the response area across all contrasts (RA; right). Red vertical lines indicate the observed accuracy of the model (peak contrast = 82.6%, RA = 84.3%). The null mean (~63%) exceeds the nominal chance level of 50% because shuffling preserves the marginal class proportions within each day, allowing incidental agreement between predicted and observed labels. This test confirms that the model's accuracy depends on the spatial and orientation structure of the data, not merely on the overall proportion of enhanced and suppressed units.

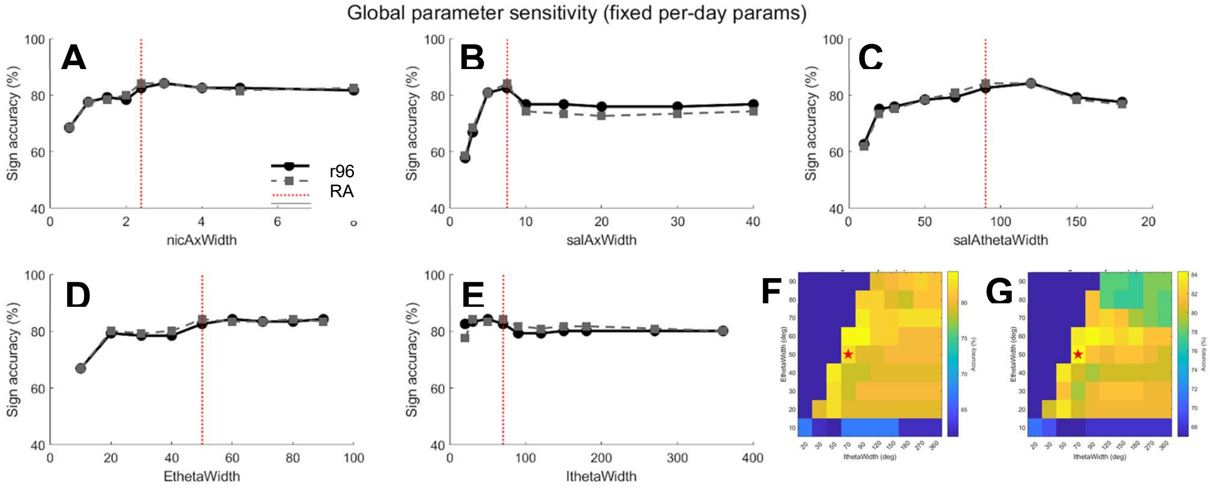


**Supplemental Figure S4. Model accuracy is robust to global parameter values.** Sign prediction accuracy was evaluated across a range of values for each shared parameter, with per-day drug parameters held fixed at their fitted values. **(A)** Nicotine spatial width (nicAxWidth). **(B)** Saline spatial width (salAxWidth). **(C)** Saline orientation width (salAθWidth). **(D)** Excitatory orientation tuning width (EθWidth). **(E)** Inhibitory orientation tuning width (IθWidth). In each panel, the red vertical line marks the value used in reported analyses. Accuracy is stable across a broad range for all parameters, with the sharpest sensitivity to salAxWidth, where very small values (< 4) degrade performance. **(F, G)** Joint sensitivity of excitatory (EθWidth; y-axis) and inhibitory (IθWidth; x-axis) model unit orientation tuning widths, shown as heatmaps for the peak contrast response (F) and response area (G) accuracy. Only the physiologically motivated region where inhibitory tuning is broader than excitatory (I > E) is shown. The red star marks fit values used (EθWidth = 50°, IθWidth = 70°). Accuracy is highest along a diagonal ridge where both widths are moderate, and degrades primarily when excitatory tuning is very narrow (< 20°). All evaluations used the same 121 extragranular units across 8 recording days. Baseline accuracy: peak contrast = 82.6%, response area = 84.3%.

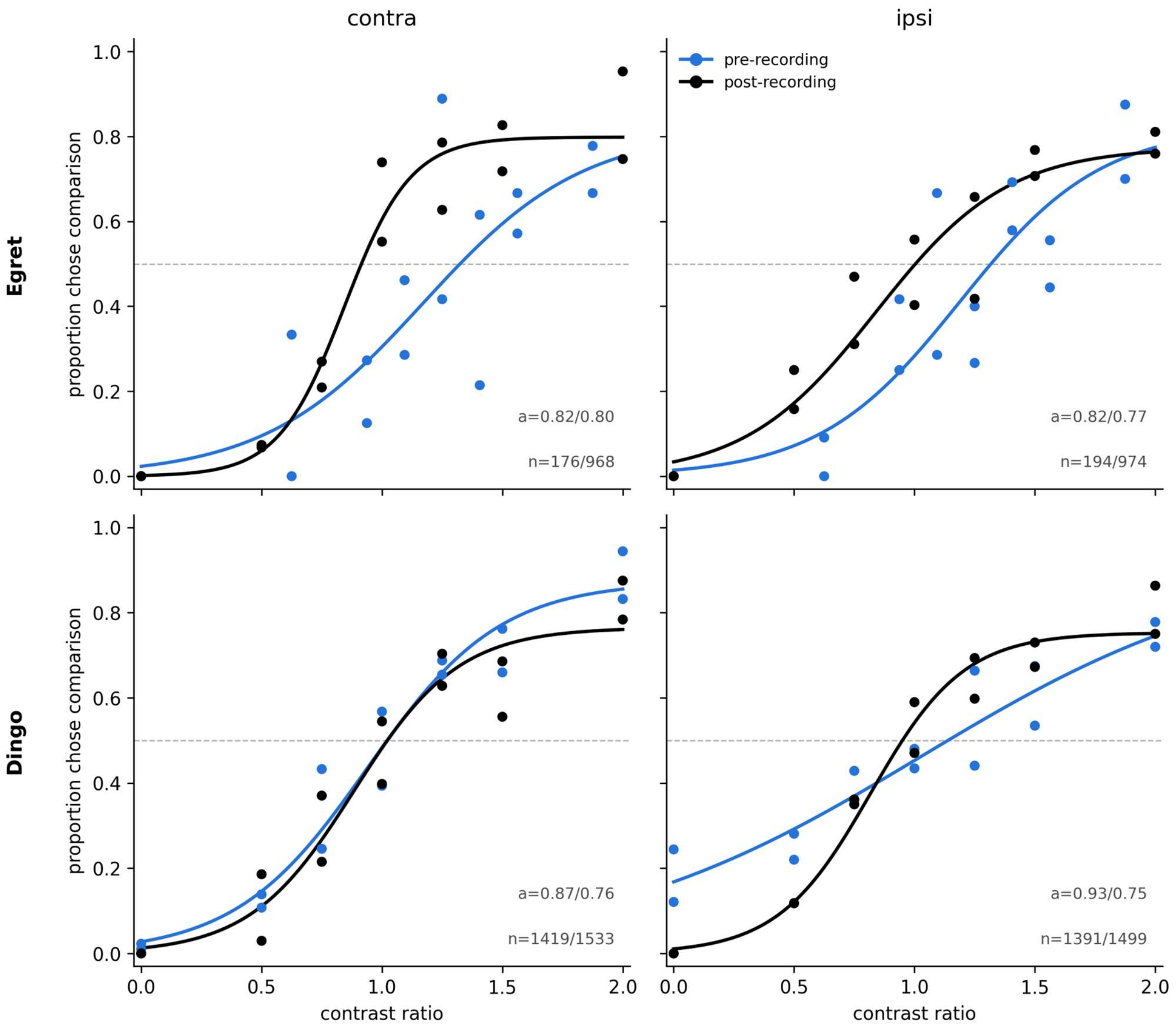


**Supplemental Figure S5. Choice behavior is comparable between hemifields on pre- and post-recording behavior-only testing days.** Choice behavior in the contralateral (left) and ipsilateral (right) hemifields is comparable in both animals E (upper) and D (lower) on behavior-only training days run both before (blue) and after (black) n vivo recording. Filled circles: session mean. Curve: logistic fits.

**Supplementary Table 1: Global model parameters.**

| Parameter | Value | Source* |
|---|---|---|
| ExWidth | 5 | Model convention |
| IxWidth | 20 | Model convention |
| EθWidth | 50 | Fit |
| IθWidth | 70 | Fit |
| Nicotine AxWidth | 2.4 | Fit |
| Saline Apeak | 4 | Model convention |
| Saline Abase | 1 | Model convention |
| Ashape | oval | Model convention |
| sigma | 1e-6 | Model convention |
| baselineMod | 0 | Model convention |
| baselineUnmod | 0 | Model convention |
| Saline AxWidth | 7.5 | Fit |
| Saline AθWidth | 90 | Fit |

*Parameters set by ‘model convention’ are unchanged from the implementation described in Reynolds and Heeger[31].
